# Molecular quantification of *Plasmodium falciparum* ookinete load reveals associations with early immune responses in *Anopheles coluzzii* mosquitoes

**DOI:** 10.64898/2026.09.21.753172

**Authors:** Giulia Bevivino, Maria Greta Dipaola, David Modiano, Bruno Arcà, Fabrizio Lombardo

## Abstract

The early stages of *Plasmodium falciparum* development in the mosquito are critical for parasite establishment and transmission, yet accurately quantifying parasite burden during ookinete maturation and midgut invasion remains challenging. Molecular markers of ookinete development offer an opportunity to investigate early infection dynamics and their association with mosquito immune responses at the individual mosquito level.

In this study, we established a molecular approach to quantify ookinete load in individual *Anopheles coluzzii* mosquitoes during the first 36 hours post-infection (hpi). Mosquitoes were fed on *P. falciparum*-infected blood with identical gametocytemia, minimizing confounding factors such as mosquito age, microbiota composition, blood meal variability, and experimental noise.

Parasite load was assessed at 12, 18, 24, and 36 hpi through the quantitative analysis of four ookinete-associated transcripts (ctrp, warp, soap, and cht1). The four markers showed a significant correlation among their transcriptional patterns during the parasite development. Moreover, the relationship between molecular marker abundance and microscopy-based oocyst counts was investigated to assess the biological relevance of this approach as an indicator of parasite infection intensity. Molecular quantification of ookinete markers revealed the presence of individual mosquitoes harboring low- and high-ookinete loads. RNA-seq analysis was performed to compare the transcriptional profiles of mosquitoes with low and high infection intensities, aiming to explore the relationship between early *P. falciparum* development and mosquito transcriptional responses. RNA-seq analysis highlighted distinct transcriptional profiles between the two groups, with mosquitoes carrying lower parasite burdens exhibiting stronger and more coordinated expression of immune-related genes, particularly in the early responses at 12 and 24 hpi. Notably, genes involved in L-arginine homeostasis were significantly modulated, highlighting a potentially relevant pathway in mosquito anti-*Plasmodium* responses.

Overall, this study highlights the utility of ookinete-associated molecular markers for quantifying *P. falciparum* infection during its earliest stages in the mosquito and linking parasite burden to vector transcriptional responses. This approach provides a valuable tool for investigating early host-vector-parasite interactions and may facilitate the identification of factors influencing parasite establishment in both laboratory and field settings.

## INTRODUCTION

Despite sustained efforts in control and prevention, malaria remains one of the most devastating parasitic diseases worldwide. According to the World Health Organization (WHO), an estimated 282 million malaria cases and over 610,000 deaths occurred globally in 2024, representing a major public health burden, particularly in sub-Saharan Africa, where the highest morbidity and mortality are reported (WHO, 2025). Malaria transmission begins when gametocytes, the sexual stages of the parasite circulating in human blood, are ingested by mosquitoes during a blood meal. The density of gametocytes in the bloodstream of infected individuals plays a crucial role in shaping malaria transmission dynamics. Human infectiousness depends on the density of both asexual parasites and gametocytes, with the latter directly influencing the probability of mosquito infection (Robert et al., 1996). However, the association between higher gametocyte densities and increased mosquito infection rates, although generally valid (Ouédraogo et al., 2009) it is not absolute, and low gametocyte densities do not necessarily preclude transmission (Schneider et al., 2007; Meibalan and Marti, 2017). In fact, membrane feeding assays, combined with sensitive molecular detection techniques, highlighted that submicroscopic gametocyte densities (<1 gametocyte/µl) may be enough to infect mosquitoes (Schneider et al., 2007). Given that the mosquito blood meal volume is approximately 2–3 µl, the presence of at least one male and one female gametocyte may be sufficient to establish infection. In this context, since parasite’s development in the mosquito requires fertilization between female and male gametes, the gametocyte sex ratio, i.e. the proportion of female to male gametocytes, is a crucial factor in evaluating and predicting mosquito infectability and a key determinant of malaria transmission (Santolamazza et al., 2017; Henry et al., 2019). Male gametocytes can produce up to eight microgametes, each capable of fertilizing a female gamete. Consequently, the gametocyte sex ratio is typically female-biased, with approximately three to four female per male gametocytes, although this ratio can fluctuate during infection (Bousema and Drakeley, 2011). Conversely, when gametocyte densities are low, transmission may be constrained by a shortage of male gametocytes, potentially leading to a relative increase in male investment to maximize fertilization success (Bradley et al., 2018).

Malaria transmission is also strongly influenced by mosquito-related factors. *Anopheles* mosquitoes possess a sophisticated innate immune system that targets invading pathogens, including *Plasmodium* parasites. The interaction between *Anopheles* mosquitoes and *Plasmodium* is highly complex and involves multiple immune responses that affect parasite survival at different stages of its development in the mosquito (Dimopoulos et al., 1998; Osta et al., 2004a, 2004b; Saab et al., 2025). These responses are active both in the mosquito midgut, the organ initially invaded by the parasite, and in the hemolymph, where systemic immune defenses are mounted. During the development from the gametocyte stage to oocyst formation in the mosquito midgut, *Plasmodium* parasites experience substantial losses, leading to a significant reduction in their numbers (Graumans et al., 2020). Indeed, out of hundreds to thousands of gametocytes typically ingested during a blood meal, only approximately 50-100 develop into ookinetes, and among these, only around five successfully progress to the oocyst stage (Sinden and Billingsley, 2001). A second major bottleneck occurs during sporozoite migration, as only a small fraction of the thousands of sporozoites produced within oocysts successfully invade the salivary glands and become transmissible to humans. A significant proportion of these losses is attributable to both local and systemic mosquito immune defenses.

Parasite elimination is mediated by two main classes of immune responses, humoral and cellular (Saab et al., 2025). Humoral immune responses, which are activated when *Plasmodium* traverses the midgut epithelium and enters the hemocoel, involve a complement-like system centered on the thioester-containing protein 1 (TEP1). TEP1 binds to *Plasmodium* ookinetes and targets them for destruction via lysis or melanization, a process facilitated by cofactors such as LRIM1 and APL1C, which stabilize the TEP1 complex (Blandin et al., 2004; Fraiture et al., 2009; Povelones et al., 2009). Additional humoral defenses include the production of antimicrobial peptides (AMPs), such as gambicin that damage parasite membranes (Vizioli et al., 2001), as well as the melanization response that encapsulates and neutralizes parasites (Barillas-Mury, 2007). Nitric oxide (NO) production in the mosquito midgut contributes to limiting *Plasmodium* development through nitrosative stress and is also implicated in systemic immune signaling, including the induction of antimicrobial peptide expression in the fat body (Luckhart et al., 1998; Herrera-Ortiz et al., 2011). On the other side, cellular immune responses are mediated by hemocytes, the principal immune cells circulating in the mosquito hemolymph.

Hemocytes are a heterogeneous population of immune cells (Rodrigues et al., 2010; Smith et al., 2015; Barletta et al., 2022; Samantsidis et al., 2025) that provide rapid and targeted defense through phagocytosis, encapsulation, and nodulation (Lavine and Strand, 2002; Castillo et al., 2006; Kumar et al., 2018). Together, humoral and cellular immune mechanisms act independently or synergistically as critical barriers to *Plasmodium* development, thereby influencing malaria transmission efficiency.

In this study, we focused on the relationship between parasite density and mosquito immune responses during the early stages of the *Plasmodium* life cycle within the mosquito, when ookinetes are formed, undergo maturation within the midgut and then traverse the midgut epithelium, that is between 12 and 36 hours post-infection. Specifically, we wished to determine whether mosquito innate immune responses correlate with infection intensity at these early developmental stages, which are crucial for parasite establishment and subsequent transmission to the human host. To minimize confounding factors (e.g., mosquito age, microbiota or blood meal composition, experimental variability in controlled infection experiment) we evaluated ookinete load in individual mosquitoes exposed to the same gametocytemia by quantifying four ookinete-specific transcripts (ctrp, warp, soap, and cht1) at 12, 18, 24 and 36 hours post-infection (hpi). This experimental scheme allowed to reliably identify mosquitoes displaying “low” and “high” ookinete loads despite being fed on blood carrying the same amount of gametocytes. RNA-seq analysis of mosquitoes with different ookinete loads was performed to evaluate a possible association between effective immune activation and containment of parasite infection.

## MATERIALS AND METHODS

### Mosquito rearing and experimental infection

Infection experiments were conducted at the Vector Biology Department of the Max Planck Institute for Infection Biology (Berlin) by transnational access within the framework of the Infravec2 project. Mosquitoes were maintained under standard insectary conditions (28°C, 70– 80% relative humidity, 12:12 h light:dark photoperiod). Adult mosquitoes were provided continuous access to a 5% sucrose solution supplied on cotton swabs. Three-day-old female *Anopheles coluzzii* mosquitoes (N’Gousso strain) were experimentally infected by membrane feeding on human blood (Haema, Berlin) containing stage V gametocytes of *Plasmodium falciparum* (NF54 strain), prepared according to standard protocols (Siciliano et al., 2020a). Two parallel infection experiments were performed in two mosquito cages using artificial membrane feeders containing identical parasite densities (3,700 gametocytes/µl). Unfed mosquitoes were removed immediately after the infectious blood meal. Fed mosquitoes were collected at 12, 18, 24, and 36 hours post-infection (hpi). In addition, a baseline time point (TP0) was included.

Approximately 50 female mosquitoes per time point (T0, T12, T18, T24, and T36) were anesthetized on ice and individually transferred into 96-well plates containing 130 µl of RNAlater solution (Thermo Fisher Scientific), ensuring complete submersion. Samples were stored at 4°C for one month and subsequently transferred to −20°C until RNA extraction. Infection prevalence (percentage of infected mosquitoes) and infection intensity (median number of oocysts per midgut) were assessed by dissecting mosquito midguts at 8 days post-infection and enumerating developing oocysts by light microscopy.

### RNA isolation

Total RNA was extracted from individual mosquitoes using two different extraction methods: phenol–chloroform extraction using TRIzol Reagent (Invitrogen) and spin-column chromatography using the RNA/DNA Purification Kit (Norgen), following the manufacturers’ protocols. For each time point (T0, T12, T18, T24, and T36), a total of 50 RNA samples were isolated, with 25 samples extracted using TRIzol and 25 using the Norgen RNA/DNA Purification Kit. Purified RNA was resuspended or eluted in 25–35 µl of RNase-free water or elution buffer, respectively, and stored at −20°C until further analysis. RNA concentration and purity were assessed by measuring absorbance at 260 and 280 nm using a BioTek Synergy HT spectrophotometer equipped with a Take3 module. RNA integrity was evaluated by electrophoresis on 1% agarose gels loading RNA amounts ranging from 80 to 500 ng.

### *P. falciparum* markers selection and primers design

Four *P. falciparum* microneme-associated genes were selected as ookinete stage markers based on microarray and RNA-seq transcriptomic data available at PlasmoDB (https://plasmodb.org): *ctrp* (circumsporozoite and TRAP-related protein), *warp* (von Willebrand factor A domain–related protein), *soap* (secreted ookinete adhesive protein), and *cht1* (chitinase 1) (Dessens et al., 1999, 2003; Yuda et al., 2001; Li et al., 2004; Ramakrishnan et al., 2011). Selection was based on the high level of expression at the ookinete stage, with increased transcript abundance during the first 24 hours post- infection (corresponding to the invasive ookinete phase) and absence or very low expression in other *Plasmodium* sexual stages (gametocytes) and mosquito-stage parasites (oocysts and sporozoites) (Abraham et al., 2004; Pradel, 2007; López-Barragán et al., 2011; Akinosoglou et al., 2015; Gómez-Díaz et al., 2017). Primers for *ctrp*, *warp*, *soap*, and *cht1* were designed (Table 1) using Primer3 software (online version: https://primer3.ut.ee/). Primer specificity was verified by BLASTn against genomic and RNA reference datasets of *P. falciparum* (PlasmoDB), *Homo sapiens* (NCBI), and *An. coluzzii* Ngousso strain (VectorBase). Primer efficiency and specificity were experimentally validated by endpoint PCR using genomic DNA from *P. falciparum* 3D7 and Platinum Taq High Fidelity DNA polymerase (Life Technologies). Each primer pair yielded a single specific amplicon of the expected size, confirming target specificity. For each gene, two primer pairs were designed: one for qPCR quantification and the other for generation of RNA standard curves. For standard curve construction, the forward primer included the T7 promoter sequence 5ʹ-TAATACGACTCACTATAG-3ʹ. A complete list of primer sequences is provided in Table 1.

**Table 1.**
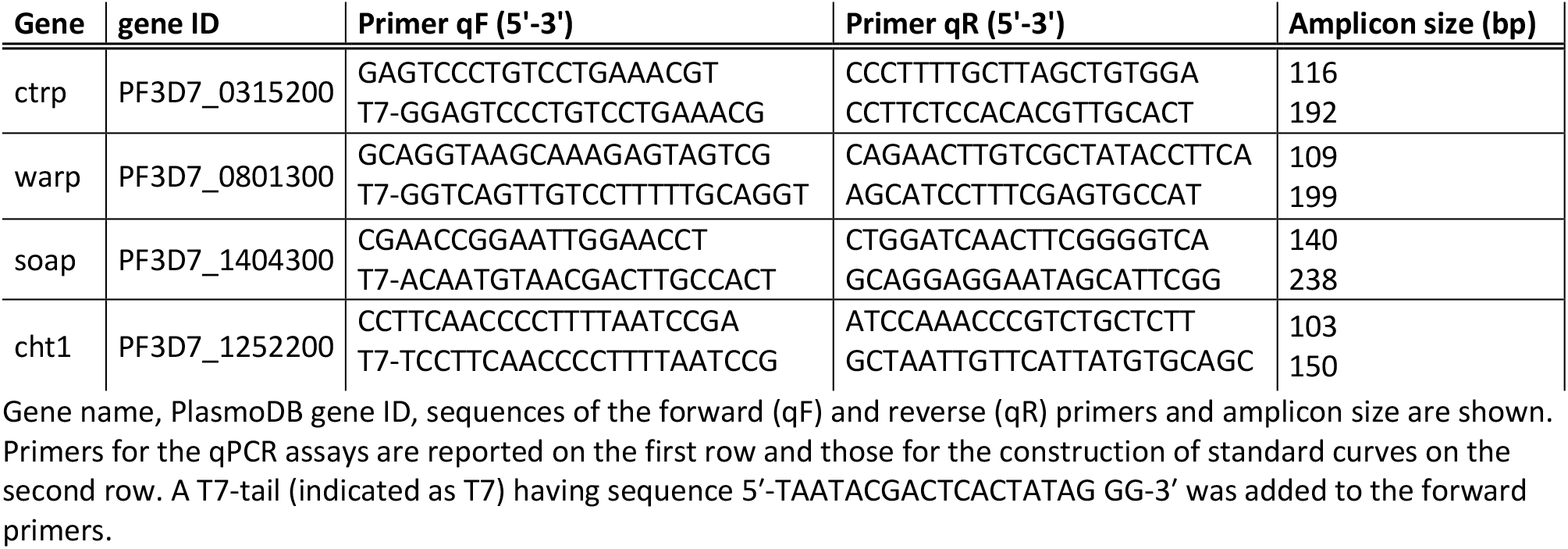
List of *P. falciparum* genes and primers for qPCR analysis.

### External standard curves

DNA amplicons for each *P. falciparum* target gene (*soap, warp, ctrp,* and *cht1*) were generated by endpoint PCR using *P. falciparum* 3D7 genomic DNA as template (Santolamazza et al., 2017), the gene-specific primers shown in Table 1 and Platinum Taq High Fidelity DNA polymerase (Life Technologies). PCR products were purified using the PureLink DNA Purification Kit (Thermo Fisher Scientific) according to the manufacturer’s instructions. Concentration and purity of the purified T7-tailed amplicons were assessed by spectrophotometry (BioTek Synergy HT, Take3 module) and their identity confirmed by Sanger sequencing. For each gene, 100 ng of purified PCR product was used as template for in vitro transcription to generate single-stranded RNA using a T7 RNA polymerase kit (Promega), following the manufacturer’s protocol. Input DNA was removed by treatment at 37°C for 15 minutes with 1 µl (2U) of TURBO DNase I (Ambion).

Synthetic RNA transcripts were purified by phenol:chloroform extraction and RNA quantity and integrity were evaluated by spectrophotometry (BioTek Synergy HT, Take3 module) and agarose gel electrophoresis. Purified synthetic RNA transcripts were reverse-transcribed using the High- Capacity RNA-to-cDNA Kit (Thermo Fisher Scientific). Transcript copy numbers were calculated using the following formula:

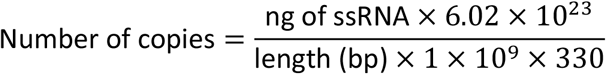

(as described at https://scienceprimer.com/copy-number-calculator-for-realtime-pcr).

The resulting cDNA was serially diluted in ten-fold steps (1:10) to generate external standard curves, with two technical replicates per dilution. Quantitative PCR efficiency (E) was calculated using the equation:

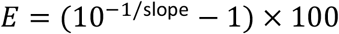

Standard curves with correlation coefficients (R²) between 0.98 and 1.00 were considered acceptable (Dhanasekaran et al., 2010).

### Real time qPCR analysis

Total RNA (1–2 µg) extracted from individual mosquitoes was DNase I-treated (Ambion TURBO DNA-free Kit) to remove contaminating genomic DNA. The efficiency of DNase I treatment was verified by endpoint PCR amplification of the *An. gambiae* AGAP004883 gene using the BIOTAQ DNA Polymerase Kit (Bioline). Briefly, approximately 20 ng of DNase I-treated RNA was used as template in a 25 µl PCR reaction containing 2.5 µl of 10× NH₄ reaction buffer, 1.2 µl of 50 mM MgCl₂, 2.4 µl of 10 mM dNTP mix, 0.5 µl each of 10 µM forward and reverse primers, 0.4 µl of BIOTAQ DNA polymerase (5U/µl), and nuclease-free water. Thermal cycling conditions were as follows: 5 minutes at 95°C; 35 cycles of 30 seconds at 94°C, 30 seconds at 55°C, and 1 minute at 72°C; followed by a final extension at 72°C for 10 minutes. Absence of amplification products was evaluated by agarose gel electrophoresis. DNase I-treated RNA (200 ng) was reverse-transcribed into cDNA using the High-Capacity cDNA Reverse Transcription Kit (Life Technologies), according to the manufacturer’s instructions. Absolute quantification was performed to determine *P. falciparum* transcript copy numbers in individual infected mosquitoes, using external standard curves generated from synthetic transcripts as described above. For each reaction, 20 ng of sample cDNA was combined with 10 µl of 2× PowerUp SYBR Green Master Mix (Thermo Fisher Scientific) and 4 µl of gene-specific forward and reverse primers (final concentration 1 µM each), in a final volume of 20 µl. Each qPCR plate included five 10-fold serial dilutions of the standard curves, expressed as transcript copy number per ng of total RNA (Devonshire et al., 2010). For each dilution point, 2 µl of synthetic cDNA was used, corresponding to initial concentrations ranging from 10⁸ to 10³ copies/ng of RNA.

Quantitative PCR was performed under the following conditions: 10 minutes at 95°C, followed by 40 cycles of 15 seconds at 95°C and 1 minute at 60°C. No-template controls (NTCs) and uninfected (TP0) mosquito samples were included in all runs to monitor contamination and baseline signal.

A total of 220 mosquitoes (44 per time point) were analysed for the abundance of ookinete- specific transcripts (ctrp, soap, warp, and cht1). Transcript copy numbers were extrapolated from the standard curves using the equation:

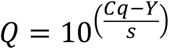

Where *Q* is the quantity of transcripts, *Cq* is the quantification cycle, *Y* is the intercept, and *s* is the slope of the standard curve. To account for differences in RNA input between samples, target transcript copy numbers were subsequently normalized to the nanograms of transcript encoding for the *An. gambiae* ribosomal protein S7 (RpS7, AGAP010592).

### RNA-seq and library preparation

Mosquitoes were classified according to *P. falciparum* infection intensity (high or low) based on the consistency of qPCR results obtained for the four ookinete-specific markers analyzed (ctrp, soap, warp, and cht1). Infection intensity was determined by quantifying the number of ookinete transcripts detected in individual mosquitoes.

Mosquitoes were assigned to the high- or low-infection-intensity groups only when concordant transcript levels were observed for at least three of the four markers. For RNA- seq analysis, three mosquitoes per condition (condition 1: high infection intensity; condition 2: low infection intensity) and per time point (T12, T24, and T36 post-infection) were selected from the qPCR dataset. RNA samples from the selected mosquitoes were pooled, resulting in one RNA pool per condition and time point. Each RNA pool was generated by combining 1 µg of total RNA from three individual infected *An. coluzzii* females, yielding a total of 3 µg of RNA per pool. RNA quality was assessed again by agarose gel electrophoresis, and residual genomic DNA contamination was removed by DNase I treatment using the TURBO DNA-free kit, as described above. Following DNase I treatment, RNA concentration was quantified using a Qubit 3.0 Fluorometer with the RNA Broad Range (BR) Assay Kit, according to the manufacturer’s instructions. RNA integrity was further evaluated using a Fragment Analyzer (Bioanalyzer) to determine the RNA Integrity Number (RIN). RNA-seq libraries were prepared following the Illumina Stranded mRNA Prep Kit protocol (IDT for Illumina UD Indexes). Briefly, polyadenylated messenger RNAs (mRNAs) were enriched using oligo(dT) magnetic beads to remove ribosomal RNA. Enriched mRNAs were then fragmented, and first- and second-strand cDNA synthesis was performed. The resulting blunt- ended cDNA fragments were adenylated at the 3ʹ ends, followed by ligation of adapters containing a 3ʹ thymine overhang. Pre-index anchor sequences were added to the cDNA fragments to enable dual indexing. Libraries were subsequently amplified by PCR to incorporate unique dual index (UDI) sequences. The UDI adapter sequences used were as follows: index 1 (i7) adapter (CAAGCAGAAGACGGCATACGAGAT[i7]GTCTCGTGGGCTCGG) and index 2 (i5) adapter (AATGATACGGCGACCACCGAGATCTACAC[i5]TCGTCGGCAGCGTC). Final library concentration was determined using a Qubit 3.0 Fluorometer, and average library size was assessed using a Bioanalyzer. Sequencing was performed on an Illumina NovaSeq 6000 platform using paired-end 2 × 150 bp reads, with a sequencing depth of approximately 30–50 million reads per library (at BMR Genomics, Padua, Italy).

### RNA-seq data processing and differential expression analysis

Raw reads were trimmed of 3ʹ adapters according to Illumina instructions and filtered (minimum length 40, minimum Phred quality score 20) using Cutadapt v4.3 (Martin and Wang, 2011). Post-trimmed quality control was performed using FastQC v0.12.1 (Andrews, S., https://www.bioinformatics.babraham.ac.uk/projects/fastqc/) and MultiQC 1.19 (Ewels et al., 2016). High-quality reads were aligned to both the *An. gambiae* PEST reference genome (AgamP4.14, GCA_000005575.2; VectorBase: https://vectorbase.org/vectorbase/app/record/dataset/NCBITAXON_180454) (Holt et al., 2002) and to the *P. falciparum* 3D7 genome (GCA_000002765.3; PlasmoDB: https://plasmodb.org/plasmo/app/record/dataset/NCBITAXON_36329) (Böhme et al., 2019) using STAR v2.7.0 (Dobin et al., 2013). Alignment was performed using species-specific intron size constraints: 20–200,000 bp for *An. gambiae* and 5–2,000 bp for *P. falciparum*. The *An. gambiae* PEST and *P. falciparum* 3D7 reference genomes were selected for sequence alignments because they represent the most comprehensively annotated genome assemblies currently available for their respective species. Gene expression quantification was carried out using RSEM (RNA-Seq by Expectation-Maximization) (Li and Dewey, 2011), and read counts were obtained separately for *An. gambiae* and *P. falciparum*. Expression levels were reported as TPM (Transcripts Per Million), which normalizes read counts by transcript length and sequencing depth. Replicate consistency was evaluated through correlation analysis, principal component analysis (PCA), and sample clustering using TrinityRNASeq v2.15.1 (Haas et al., 2013). Differential expression analysis was performed using edgeR. Pairwise comparisons were conducted between mosquitoes with high and low infection intensity at each time point: 12, 24, and 36 hpi. Statistical significance was assessed by adjusting p-values using the false discovery rate (FDR) method. Only genes with an adjusted p- value ≤ 0.05 and an absolute log₂ fold change ≥ 1 were considered differentially expressed.

Functional annotation of differentially expressed genes, including InterPro IDs, Pfam IDs, and Pfam descriptions, was retrieved from the VectorBase Research database. Gene Ontology (GO) enrichment analysis was performed using the PANTHER classification system (https://pantherdb.org/) (Mi et al., 2017). Enrichment analyses were conducted for the three main GO categories, i.e., molecular function, biological process, and cellular component, using differentially expressed genes identified by edgeR and the complete set of expressed genes at each time point (12, 24, and 36 hpi) as reference. Statistical significance was evaluated using Fisher’s exact test, with a significance threshold of p < 0.05 and a fold enrichment (FE) > 2. FASTQ files containing RNA-Seq data are deposited in the NCBI SRA database under bioproject accession numbers GSE337583.

### Validation of RNA-seq results by qPCR

Primers targeting the mosquito genes *SCRBQ2*, *ANXB9*, *LRR*, *Peritrophin-A*, *Carboxypeptidase B*, *argininosuccinate synthase (ASS)*, and *Ficolin-A* were designed using Primer3 (https://primer3.ut.ee/). Primer specificity was evaluated by BLASTn against *P. falciparum* (PlasmoDB), *H. sapiens* (NCBI), and *An. coluzzii* Ngousso strain (VectorBase) reference sequences to exclude potential cross-amplification. Primer sequences and gene identifiers are listed in Table 2. Total RNA was extracted from four *An. coluzzii* adult females (N’gousso strain) 2-5-day-old using the TRIzol reagent. RNA quality and concentration were assessed spectrophotometrically (BioTek SynergyHT, Take3 module), and RNA integrity was verified by agarose gel electrophoresis. Following DNase I treatment, cDNA was synthesized as described previously. Standard curves were generated from serial dilutions of cDNA (100, 20, 4, 0.8, and 0.016 ng/µL). Relative gene expression was measured by qPCR using cDNA pools prepared from individual mosquitoes (three mosquitoes per pool). Reactions were performed in 20 µL containing 2 µL cDNA (1 ng/µL), 10 µL PowerUp SYBR Green Master Mix (2X), and 200 nM of each primer. Amplification was carried out with an initial denaturation at 95°C for 10 min, followed by 40 cycles of 95°C for 15 s and 60°C for 1 min. Melting curve analysis was performed to confirm amplification specificity. All samples were analyzed in duplicate, and no-template controls (NTC) were included in each run. Gene expression was normalized using the endogenous reference gene RpS7 (AGAP010592), encoding the ribosomal protein S7. The T12 time point was used as the calibrator. Relative expression levels were calculated using both the standard curve method and the ΔΔCT method. For ΔΔCT analysis, ΔCT values were calculated by subtracting RpS7 Ct values from target gene Ct values, and fold changes were expressed as 2⁻^ΔΔCT^. When sufficient biological replicates were available, statistical differences between groups were evaluated using Student’s t-test, followed by the Wilcoxon test. Graphs were generated using GraphPad Prism 9.

**Table 2.**
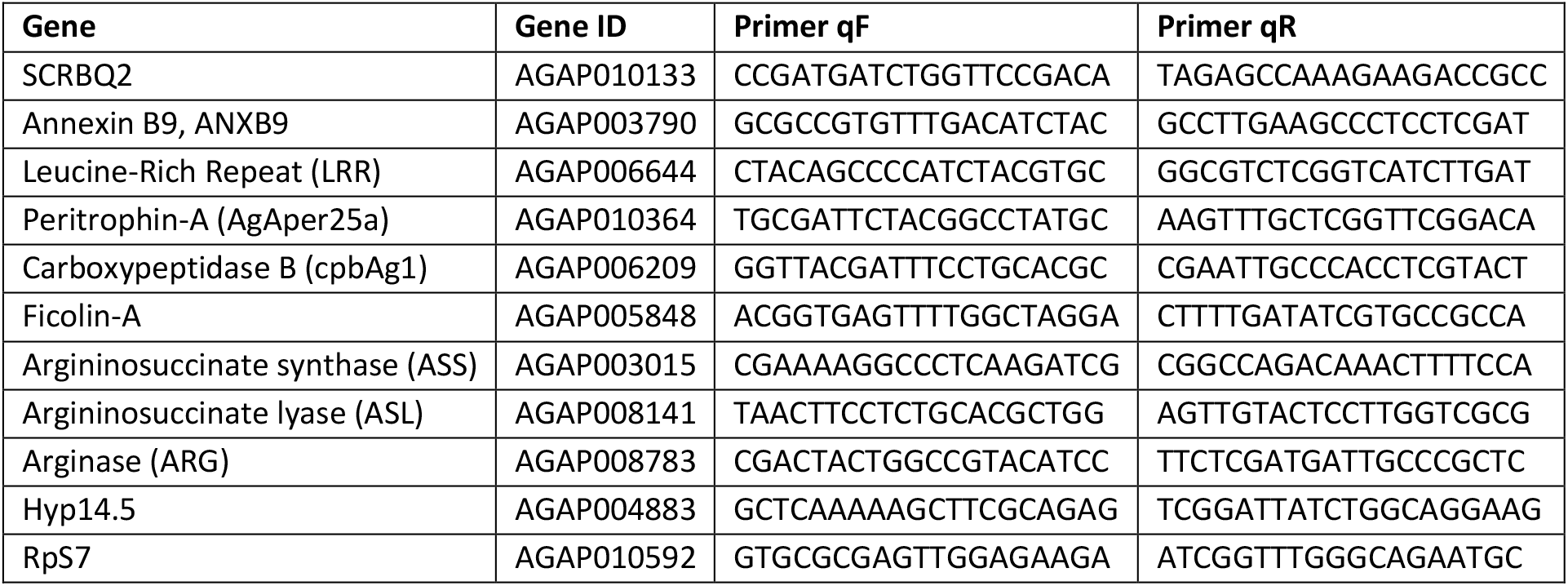
List of *An. gambiae* primers used for qPCR validation.

## RESULTS

### Mosquito infection

Artificial membrane feeding remains the gold standard method for studying *P. falciparum* transmission from humans to mosquitoes and for assessing mosquito susceptibility and infectivity. We used membrane feeding assays to infect *An. coluzzii* mosquitoes with *P. falciparum* NF54 (Figure 1A). Mosquitoes were collected at 12, 18, 24, and 36 hours post-infection (hpi), corresponding respectively to early, mature, invasive, and late ookinete stages. Unfed mosquitoes (TP0) were removed immediately after feeding.

**Figure 1.**
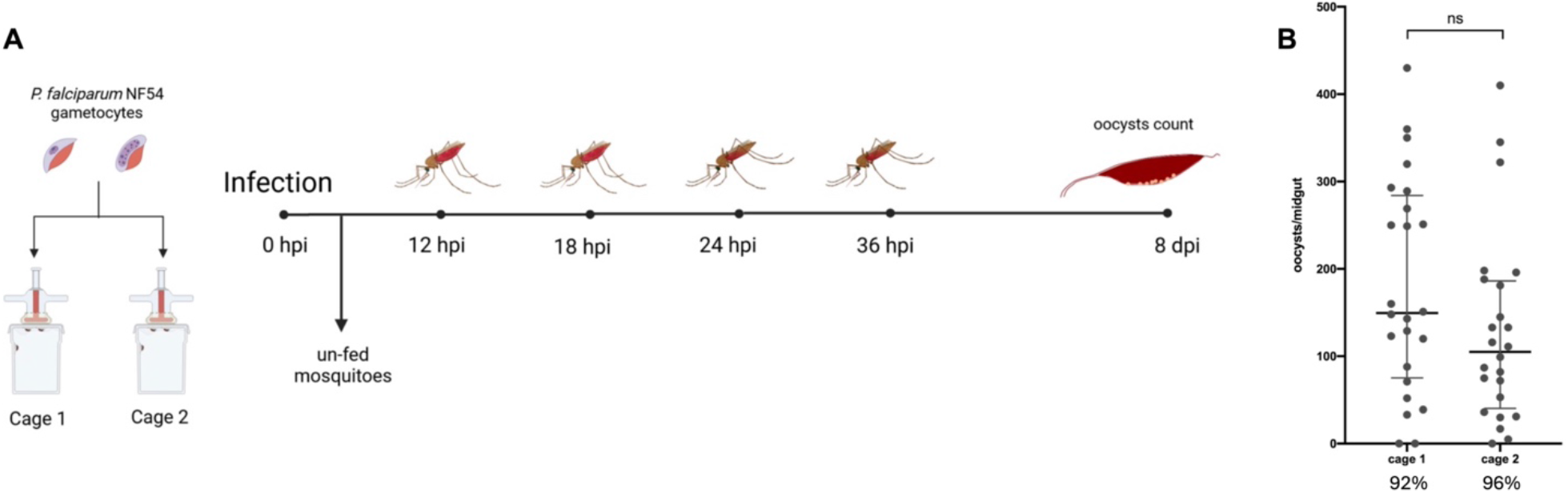
**Overview of the experimental infection with *P. falciparum* and mosquito collection**. A. Two cages of *An. coluzzii* mosquitoes from the same generation were used for parallel infection with *P. falciparum* NF54 stage V gametocytes at a concentration of 3700 gametocytes/µl. Mosquitoes were singularly collected at 0 (non-infected), 12, 18, 24, and 36 hours after infection. The figure also shows blood meal digestion in the mosquito midgut over the first 36 hours after infection. B. The intensity of infection was evaluated by oocyst count in the midguts of 24 individual mosquitoes per cage at 8 days post-infection. The number of oocysts per midgut in each cage are shown by the grey little dots. Horizontal lines denote median oocyst counts and error bars the interquartile range. Below x-axis are reported the infection prevalences for each cage. Statistical comparison between groups was performed using the Mann–Whitney test.

Infections were conducted in parallel using two cages (cage 1 and cage 2), each containing 150–200 females from the same generation, which were fed identical aliquots of human blood containing 3700 gametocytes/µl. At each time point, at least 50 mosquitoes (approximately 25 per cage) were collected. Infection outcome was assessed by counting oocysts in midguts dissected 8 days post-feeding, with a total of 48 mosquitoes (24 per cage) analyzed. Median oocyst numbers and infection prevalence were slightly different between cage 1 (149.5 and 92%, respectively) and cage 2 (105 and 96%), but this difference was not statistically significant (Figure 1B).

### Transcriptional profiling and absolute quantification of the ookinete marker transcripts

Four *P. falciparum* ookinete-specific genes (*ctrp*, *warp*, *soap*, and *cht1*; Table 3), with minimal expression in other parasite stages according to published RNA-seq, microarray, and single-cell transcriptomic datasets, were selected to reveal and quantify ookinete development in individual mosquitoes (López-Barragán et al., 2011; Akinosoglou et al., 2015; Real et al., 2021).

**Table 3.**
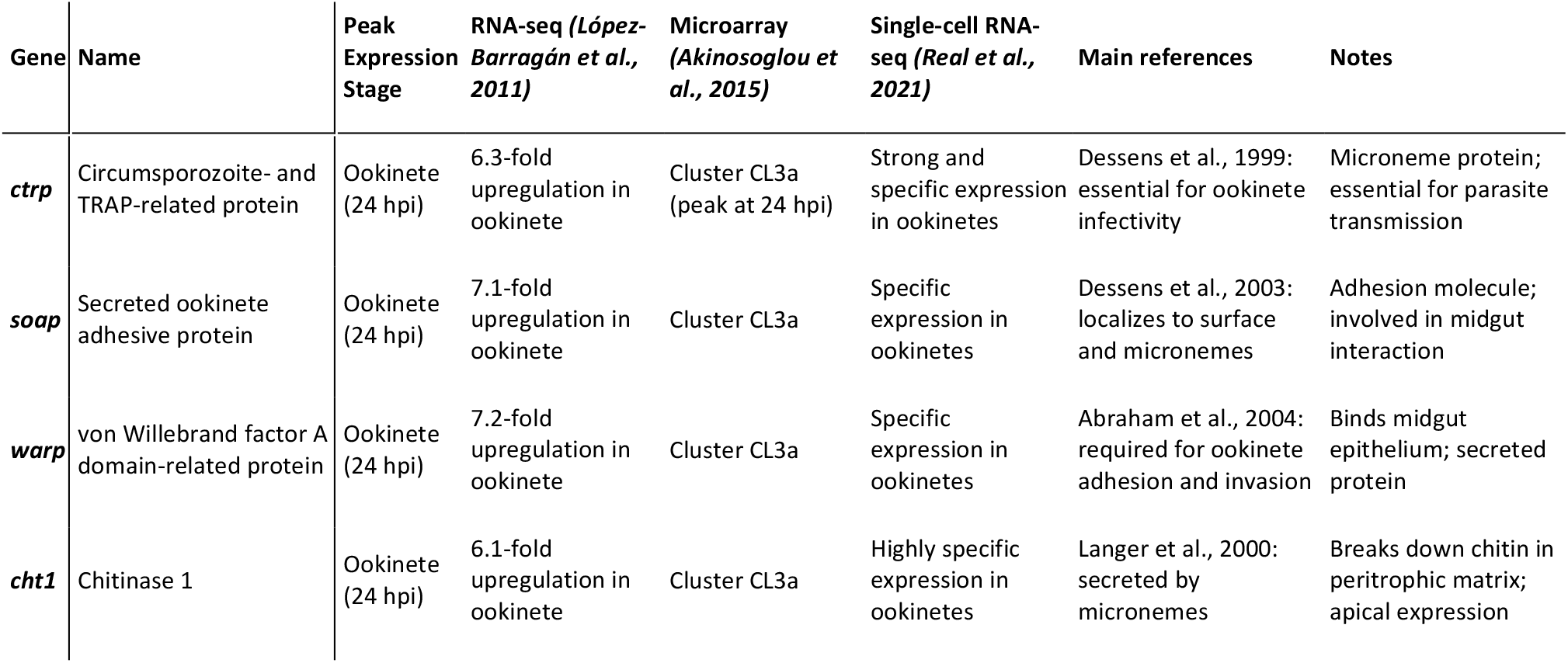
Ookinete markers candidates selected for this study: main properties and references.

To verify, in our experimental conditions, the transcriptional profiles of these ookinete markers, we initially analyzed their temporal expression patterns in pools of infected mosquitoes across the four selected time points. Each pool consisted of RNA extracted from eight randomly selected individuals, and relative quantification was applied to analyze the real-time PCR data. Results confirmed the pattern expected for the four markers, with a marked transcript increase between 12 and 18 hpi, a transcriptional peak at 24 hours, and a decline by 36 hpi, consistent with known ookinete development dynamics (Figure S1). Soap, warp, and cht1 transcripts showed the strongest induction (4–6-fold), whereas ctrp increased more modestly. Next, to assess potential inter-individual variations and provide a more robust evaluation, we refined the analysis of ookinete marker expression at the single-mosquito level across the different time points. To this end, synthetic RNAs and transcript-specific standard curves were generated to enable absolute quantification of the four ookinete markers in real-time quantitative PCR (qPCR) experiments. Standard curves were validated across eight independent experiments using ten-point serial dilutions (Supporting File S1, Figures S2 and S3). qPCR analysis was then performed on individual mosquitoes collected at 12, 18, 24, and 36 hpi. Approximately 44 *An. coluzzii* females per time point were analyzed individually using gene-specific primers, SYBR Green chemistry, and an absolute quantification approach, as described in the Materials and Methods. Transcript copy numbers for each marker gene were determined by interpolating sample Ct values against standard curves included on each qPCR plate, then normalized to RpS7 transcript abundance to account for differences in RNA input between samples (Figure 2). The transcriptional trends observed in the single-mosquito analysis resulted coherent with observations from pooled mosquito samples.

**Figure 2.**
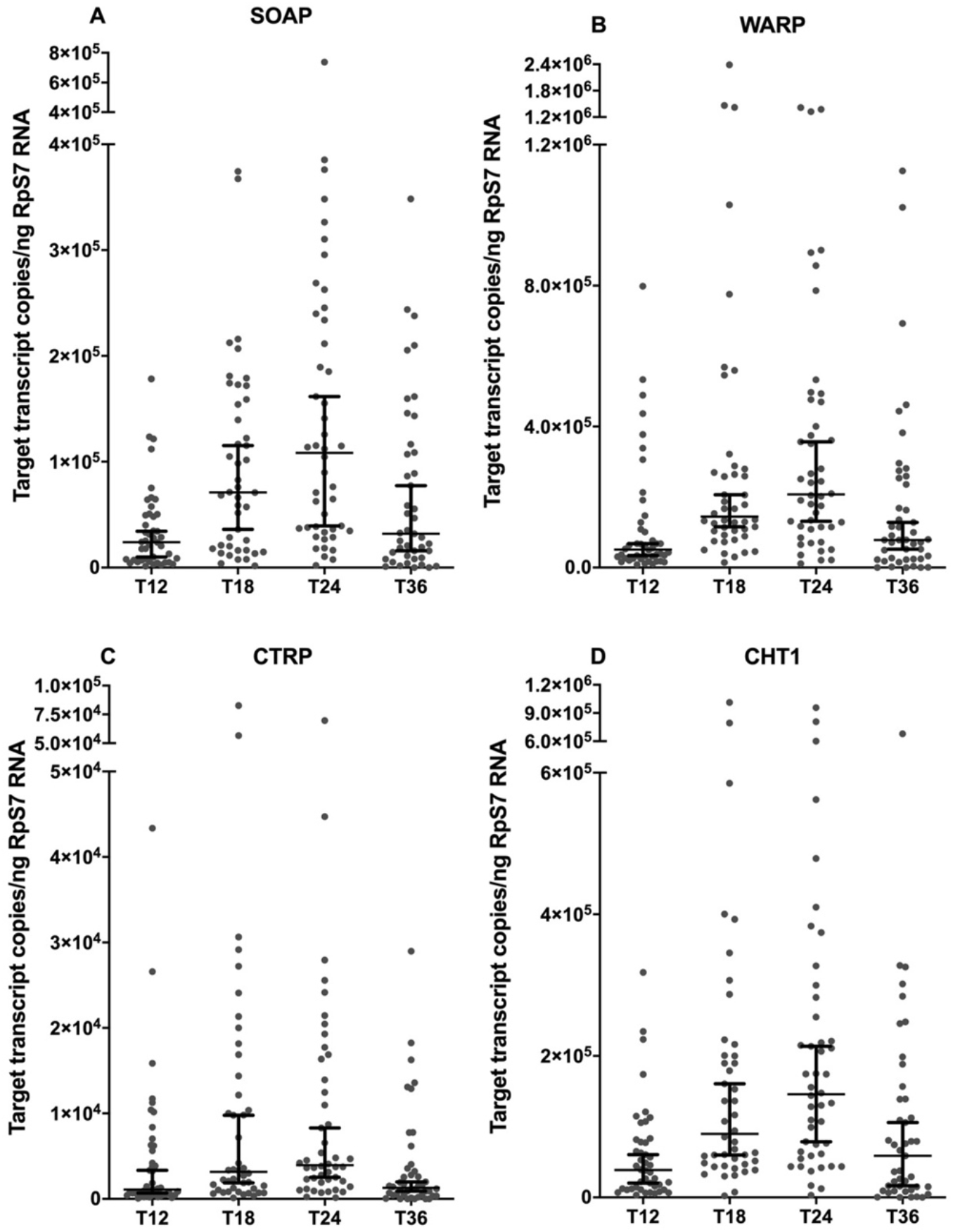
Absolute quantification of *P. falciparum* ookinete markers (ctrp, warp, soap and cht1) in single *An. coluzzii* infected mosquitoes. The different time points post infection (T12, T18, T24 and T36 hpi) and transcript copy number normalized to nanograms of the RpS7 gene are reported for each ookinete marker (A, soap; B, warp; C, ctrp and D, cht1). Horizontal lines indicate median values with 95% confidence interval. For each graph, gene expression data relative to 44 mosquitoes/time point are reported.

For all markers, transcript abundance increased over time and reached a maximum at 24 hpi, consistent with the timing of the invasive ookinete stage. As observed for oocyst counts by microscopy, substantial inter-individual variability in marker transcript abundance was detected at all time points, underscoring the pronounced heterogeneity of parasite development among mosquitoes within the same experimental cohort.

### Pairwise correlation analysis of soap, cht1, ctrp, and warp transcripts

The temporal coherence of expression among the four marker transcripts at the different time points of invasion was evaluated by Pearson’s correlation analysis (Figure 3). Cumulative analysis of the four time points (TPs) showed strong positive correlations between soap and cht1 (Figure 3A: r = 0.7681, P < 0.0001), cht1 and ctrp (Figure 3B: r = 0.7817, P < 0.0001), and ctrp and warp (Figure 3F: r = 0.8275, P < 0.0001). Moderate correlations were detected between cht1 and warp (Figure 3D) and between soap and ctrp (Figure 3C), while a weaker association was observed between soap and warp (Figure 3E). Overall, these results indicate a coordinated expression pattern among core invasion-associated markers (cht1, ctrp, and warp), with soap showing comparatively lower concordance, particularly with warp. When the different TPs were analyzed independently, correlation among the different ookinete markers could be further refined. At 12 hpi, cht1, ctrp, and warp showed strong positive correlations (Figure 3G: r = 0.57–0.90, p ≤ 0.001), suggesting early co-regulation during ookinete differentiation. In contrast, soap displayed only moderate correlations with cht1 (r = 0.54) and weaker associations with ctrp (r = 0.39) and warp (r = 0.26, not significant), indicating partial transcriptional asynchrony at this early stage. At 18 hpi, correlations strengthened overall. cht1, ctrp, and warp remained highly correlated (Figure 3H: r = 0.72–0.90, p ≤ 0.0001); soap showed increased correlation with cht1 (r = 0.64) and ctrp (r = 0.49), while remaining weakly associated with warp (r = 0.24, ns). At 24 hpi, strong correlations persisted among cht1, ctrp, and warp (Figure 3I: r = 0.67–0.74, p ≤ 0.0001), whereas the association between soap and warp was lost (r = 0.01, ns). The correlation between soap and cht1 further increased (r = 0.78, p ≤ 0.0001), suggesting closer coupling of these markers at peak ookinete development. At 36 hpi, high correlations were maintained among cht1, ctrp, and warp (Figure 3J: r = 0.70–0.83, p ≤ 0.0001), indicating sustained co-expression during late invasion/early oocyst transition. soap exhibited strong correlation with cht1 (r = 0.86) and moderate correlation with ctrp (r = 0.63) but remained weakly associated with warp (r = 0.13, ns). Overall, these data highlight a tightly coordinated transcriptional program among core ookinete invasion markers (cht1, ctrp, warp), while soap displays a partially distinct temporal regulation pattern, particularly in relation to warp. This pattern likely reflects stage-specific functional specialization during ookinete maturation and midgut traversal.

**Figure 3.**
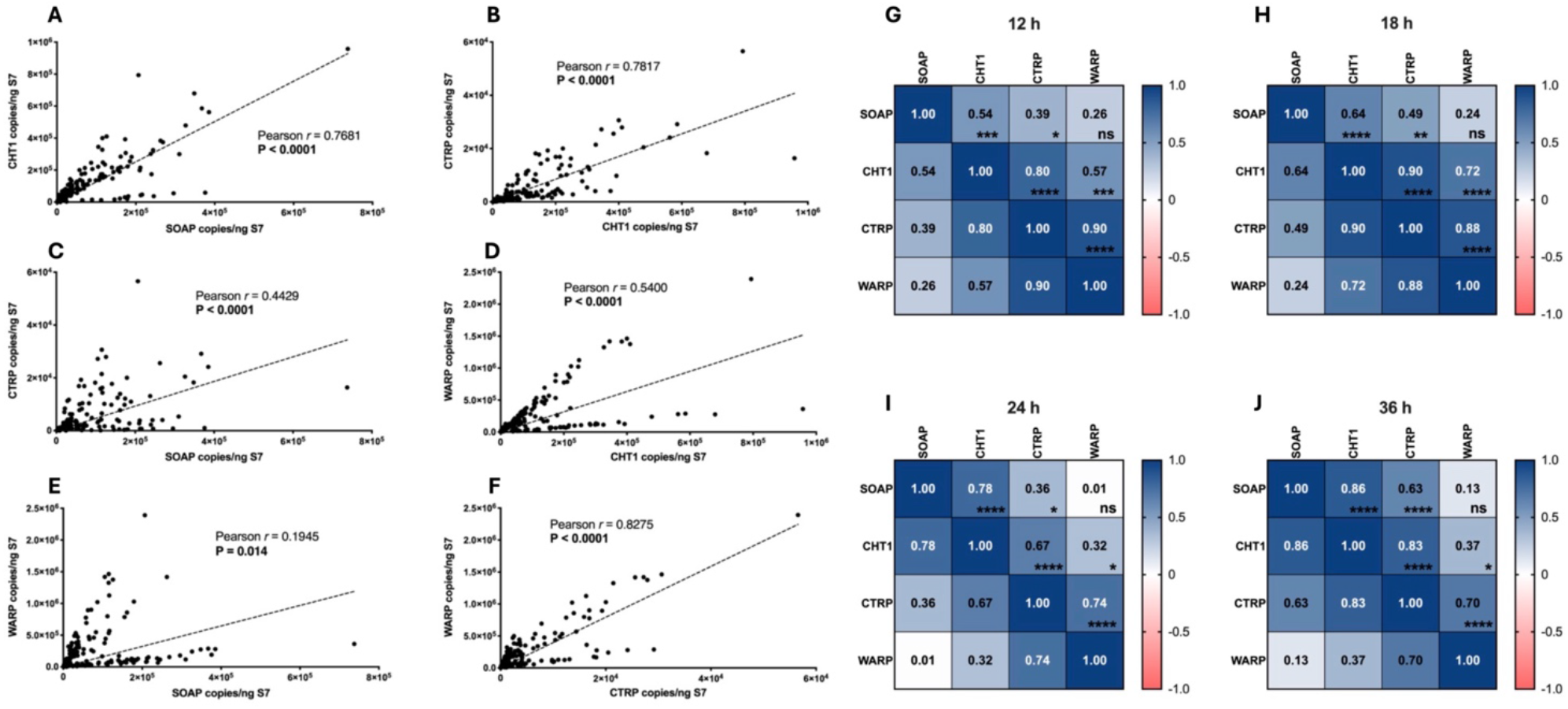
Correlation among *P. falciparum* ookinete marker transcripts across time points. (A–F) Pairwise scatter plots showing correlations between copy number (normalized per ng RpS7 RNA) of the four ookinete marker transcripts (soap, cht1, ctrp, and warp) measured by qPCR in individual mosquitoes at the four time points. Each point represents a single mosquito. Dashed lines indicate linear regression fits. Pearson correlation coefficients (r) and corresponding P values are indicated in each panel. (G–J) Heatmaps showing Pearson correlation matrices among the four ookinete markers at the four different time points post-infection: 12 hpi (G), 18 hpi (H), 24 hpi (I), and 36 hpi (J). Correlation coefficients (r) are shown within each cell and are represented by a colour scale ranging from −1 (red) to +1 (blue). Statistical significance: *, P < 0.05; **, P < 0.01; ***, P < 0.001; ****, P < 0.0001; ns, not significant.

### Correlation between ookinete marker expression and oocyst burden

As noted above (see Figure 1), oocyst counts obtained by microscopy revealed modest differences between the two mosquito cages used for experimental infections, with a higher median oocyst burden observed in mosquitoes from cage 1 compared to cage 2. To assess whether this variability was also detectable at the molecular level at the ookinete stage, we compared the transcript abundance of ookinete-specific markers between mosquitoes from the two cages at the different time points. This analysis aimed to directly relate the number of ookinete marker transcripts measured in individual mosquitoes during the late time-points of the ookinete development (i.e., 24 and 36 hpi) to the subsequent oocyst burden measured by microscopy. In principle, higher expression levels of ookinete-specific genes , especially during the late maturation of ookinetes, might correlate with higher oocyst counts at later stage. Consistent with this expectation, a similar trend was observed across all four ookinete markers at both 24 and 36 hours post infection. Mosquitoes from cage 1 (median oocyst count = 149.5) showed higher median transcript levels for all four markers than mosquitoes from cage 2 (median oocyst count = 105) (Figure 4).

**Figure 4.**
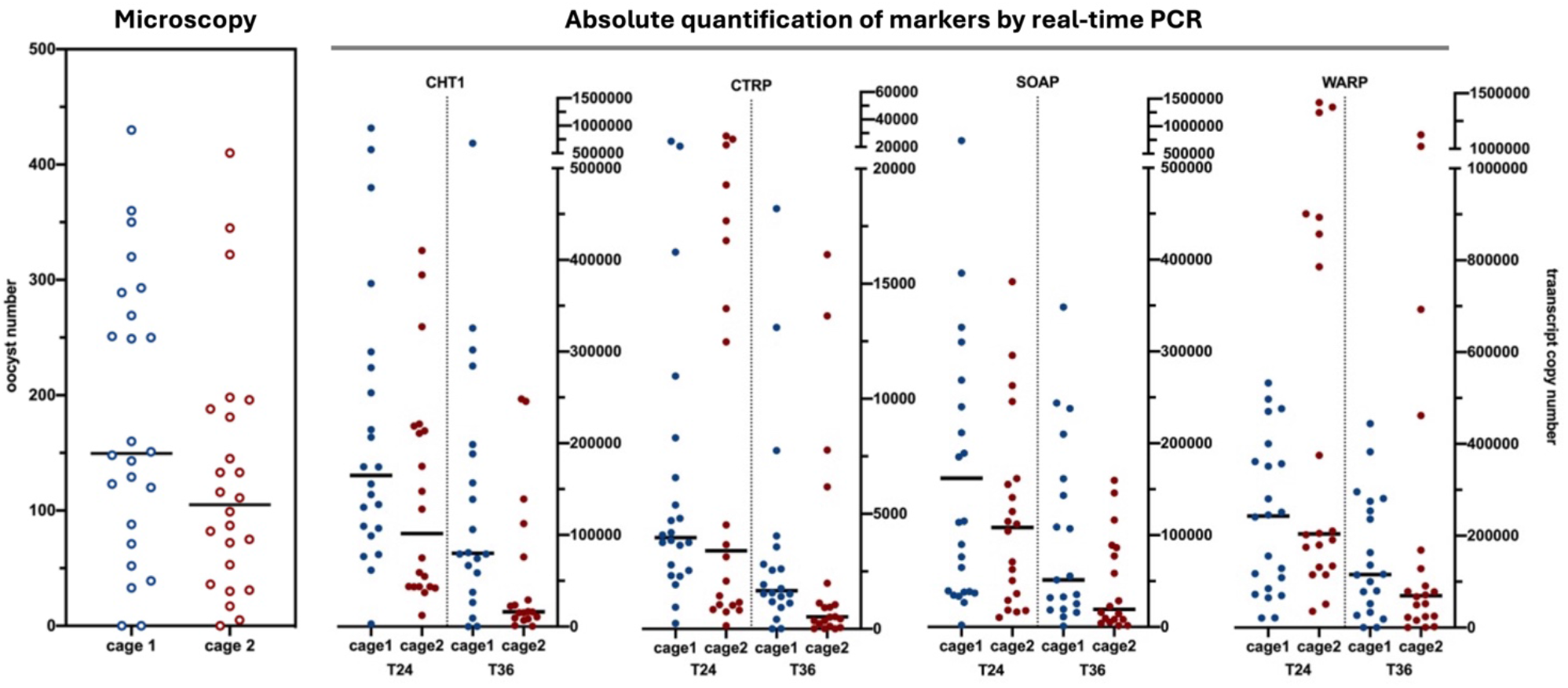
Oocyst burden and ookinete marker expression in individual mosquitoes from the two cages at 24 h and 36 h post-blood meal. Left panel: individual midgut oocyst counts in mosquitoes from cage 1 (blue) and cage 2 (red). Each dot represents a single mosquito; horizontal bars indicate the medians (data as in Figure 1B). Right panel: quantitative analysis of the ookinete marker transcripts cht1, ctrp, soap and warp at 24 h (T24) and 36 h (T36) post- blood meal. Each dot represents a single mosquito, blue from cage 1 and red from cage 2; horizontal bars indicate the medians.

Statistical analysis using two-way ANOVA followed by Tukey’s multiple comparison test did not reveal significant differences between cages for comparisons showed in Figure 4. This result mirrors the microscopy-based oocyst counts, where differences between cages were also not statistically significant, despite a higher infection burden in cage 1 (Figure 1 and Figure 4).

### Transcriptomic profiling of mosquitoes with high and low ookinete infection intensity

qPCR analysis of *P. falciparum* ookinete markers (ctrp, warp, soap, and cht1) revealed substantial inter-individual variability in transcript copy numbers among mosquitoes; this fluctuation can be assumed to reflect a variation in ookinete numbers, and a similar variation was found when considering oocysts number (see Figure 2 and 3). These observations indicate that, despite experimental variables were reduced to a minimum (mosquitoes of the same laboratory strain, same age, reared under identical environmental conditions and originated from the same infection experiment), yet individual mosquitoes displayed markedly different parasite loads following exposure to the same gametocytemia. Some variability in the volume of blood ingested by different individualsmay at least in part account for these differences. However, early mosquito responses to the infectious blood meal are known to play a key role in limiting parasite infection; therefore, distinct transcriptional responses to an identical parasite challenge mounted by individual mosquitoes may result in different levels of parasite development, even when external variables are minimized. For these reasons, we decided to analyze by RNA-seq the transcriptional responses of *An. coluzzii* mosquitoes exhibiting low versus high ookinete loads, as determined by qPCR analysis (Figure 5 and Figure S4). We focused on three key time points, 12, 24, and 36 hpi, which correspond to critical phases of parasite development in the midgut, including epithelial traversal and the first major developmental bottleneck, largely driven by the mosquito innate immune response. We chose to perform RNA-seq on pools of three mosquitoes rather than on single individuals because, despite the strong correlations observed among markers (Figure 4), a certain variability remained, and pooling samples allowed to minimize the risk of misclassification based on individual marker fluctuations. RNA was extracted from whole mosquitoes. Although this approach precluded the characterization of midgut-specific responses during early parasite development, it enabled the assessment of broader transcriptional changes potentially reflecting both local and systemic immune and metabolic responses.

**Figure 5.**
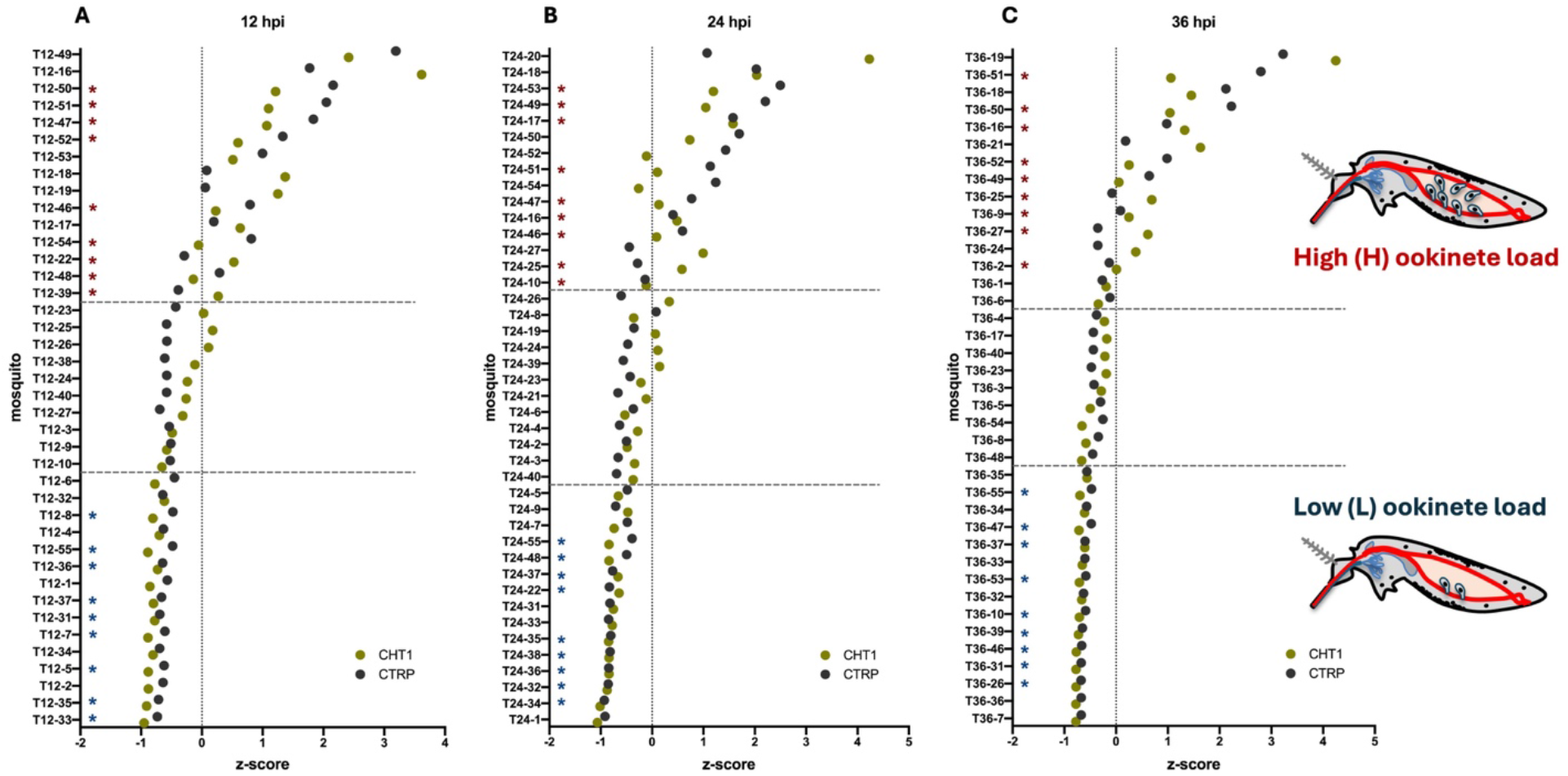
Distribution of ookinete marker expression in individual mosquitoes at different time points. Scatter plots showing z-score–normalized transcript levels of cht1 and ctrp in individual mosquitoes at 12 (A), 24 (B), and 36 (C) hpi. Each dot represents the expression level of a marker in a single mosquito. Mosquitoes are ordered along the y-axis according to increasing infection intensity, from the lowest (bottom) to the highest (top). The vertical dashed line indicates a z-score of 0 (mean expression level). Horizontal dashed lines delimit the 15 mosquitoes with the highest (H) and lowest (L) expression levels of both markers. Blue and red asterisks indicate the mosquitoes selected for RNA- seq analysis (three groups of three mosquitoes for each infection level), representing low (L) and high (H) ookinete burdens, respectively.

To generate RNA pools for sequencing, we selected mosquitoes based on the transcript levels of the cht1 and ctrp markers, as these showed the strongest correlation across most time points. Three groups of three mosquitoes were therefore chosen from the fifteen individuals exhibiting the highest marker expression (high ookinete load, H) and the fifteen exhibiting the lowest marker expression (low ookinete load, L, Figure 5). To validate this selection, the expression levels of all four ookinete markers (ctrp, soap, warp, and cht1) were also examined. This confirmed that only mosquitoes consistently displaying either "high" or "low" transcript levels for at least three of the four markers were included (Figure S4). A small number of exceptions were made when RNA quantity or quality limited the availability of suitable samples for RNA-seq analysis (Supporting File S1, Figure S4 and Supporting Excel File S1).

Total RNA from three mosquitoes with low (L) or high (H) ookinete loads at each time point (12, 24, and 36 hpi) was pooled, generating the following sample groups: T12-L, T24-L, T36-L, T12-H, T24-H, and T36-H. For each group, three independent replicates were prepared and processed for RNA-seq as described in the Materials and Methods. Details about the composition of each pool are reported in Supporting Excel File S1.

### RNA-seq sequencing quality and mapping statistics

RNA-seq libraries were prepared and sequenced following standard protocols, as detailed in the Materials and Methods section. Quality control confirmed high-quality sequencing data with a total of 664 million reads generated. Following trimming and filtering, 497.55 million reads were retained for alignment against the *An. gambiae* PEST and *P. falciparum* 3D7 reference genomes. Alignment to the *An. gambiae* reference genome yielded an overall mapping rate of 74.9% across samples (Table S1), with 13,845 contigs assembled (Supporting Excel File S2). Alignment to the *P. falciparum* 3D7 reference genome resulted in the assembly of 5,720 contigs (Supporting Excel File S2).

Correlation and cluster analysis confirmed strong consistency among biological replicates and clear segregation of samples according to experimental groups. Notably, samples clustered according to both infection intensity (high vs. low, as determined by qPCR) and time point post- infection (T12, T24, and T36), corresponding to key stages of ookinete development in the mosquito midgut (Supporting File S1, Figure S5 and S6). Together, these results confirm the robustness of the sequencing data and indicate that the experimental design effectively captured biologically meaningful transcriptional variation across conditions.

### Concordance between qPCR and RNA-seq validates infection intensity classification

To directly assess the concordance between qPCR and RNA-seq quantification, we examined the expression levels of the four ookinete markers (ctrp, warp, soap, and cht1) in the RNA-seq datasets. For each of the four markers, transcript abundance expressed as TPM (Transcripts per Million) was compared to qPCR copy numbers. Pearson correlation analyses showed a significant positive correlation for all ookinete markers, indicating strong agreement between the two quantification methods (Supporting File S1, Figure S8). These results validate our mosquito stratification strategy based on absolute quantification of ookinete markers using real time qPCR. The defined experimental groups, based on infection intensity (H vs. L) and time post- infection (T12, T24, and T36), therefore provide a robust framework for investigating how mosquito transcriptional responses vary with parasite burden during early *Plasmodium* development.

We have also analysed the Differential expression (DE) analysis identified a limited number of *Plasmodium* transcripts showing significant differences between mosquitoes with high and low infection intensity across the analyzed time points (Supporting File S1, Figure S7). At 12 hpi, ctrp and cht1, together with the important ookinete-associated gene PIMMS43 (Ukegbu et al., 2020), were significantly more abundant in mosquitoes with high infection intensity. Similarly, at 36 hpi, warp, along with the ookinete-associated genes PIMMS1, PIMMS57, and SOPT (Armistead et al., 2018; Ukegbu et al., 2021), also showed significantly higher transcript abundance in the high-infection group (Supporting Excel File S5).

### Differential expression and GO enrichment analyses

We next focused on mosquito genes, gene families and pathways that were differentially modulated across the experimental conditions. Differentially expressed (DE) genes were identified using the edgeR package by comparing mosquitoes with high (H) and low (L) infection intensity at each time point (T12, T24, and T36). Results of the differential expression analysis are summarized in Table 4.

**Table 4.**
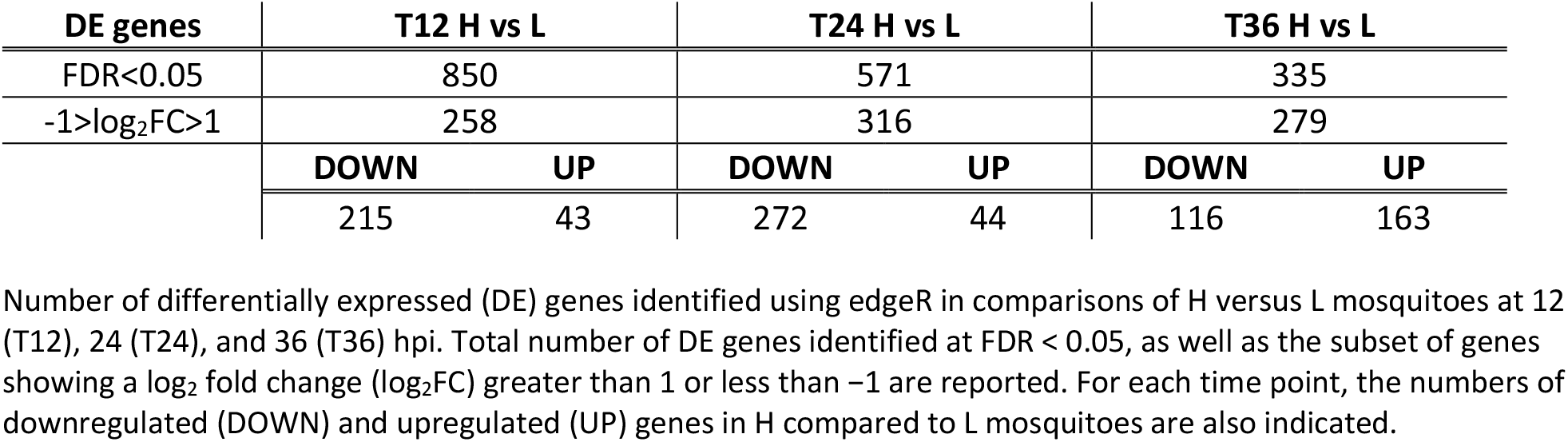
Differentially expressed genes between high (H) and low (L) infection intensity mosquitoes.

| **DE genes** | **T12 H vs L** |  | **T24 H vs L** |  | **T36 H vs L** |  |
|---|---|---|---|---|---|---|
| FDR<0.05 | 850 |  | 571 |  | 335 |  |
| -1>log<sub>2</sub>FC>1 | 258 |  | 316 |  | 279 |  |
|  | **DOWN** | **UP** | **DOWN** | **UP** | **DOWN** | **UP** |
|  | 215 | 43 | 272 | 44 | 116 | 163 |

At 12 hours post-infectious blood meal, corresponding to the early stage of ookinete development, when only a fraction of ookinetes is at the onset of midgut invasion while part of them is still in the blood bolus, differential expression analysis identified 43 genes significantly upregulated in mosquitoes with high *Plasmodium* infection intensity. In contrast, 215 genes were significantly downregulated in the high-infection group (i.e., upregulated in mosquitoes with low parasite loads) (Figure 6A). These results indicate that early differences in parasite burden are already associated with pronounced and distinct mosquito transcriptional responses during the initial phase of midgut invasion. At 24 hours post-infectious blood meal, corresponding to ookinete invasion of the midgut epithelium, 44 genes were significantly upregulated in mosquitoes with high infection intensity, whereas 272 genes were significantly downregulated in the high- infection group (i.e., upregulated in low-infection mosquitoes) (Figure 6B). This pattern suggests a sustained and dynamic host transcriptional response that continues to diverge according to parasite burden during the invasive phase. Finally, at 36 hours post-infectious blood meal, corresponding to the late ookinete stage and the onset of early oocyst formation, differential expression analysis identified 163 genes significantly upregulated and 116 genes significantly downregulated in mosquitoes with high infection intensity (Figure 6C).

**Figure 6.**
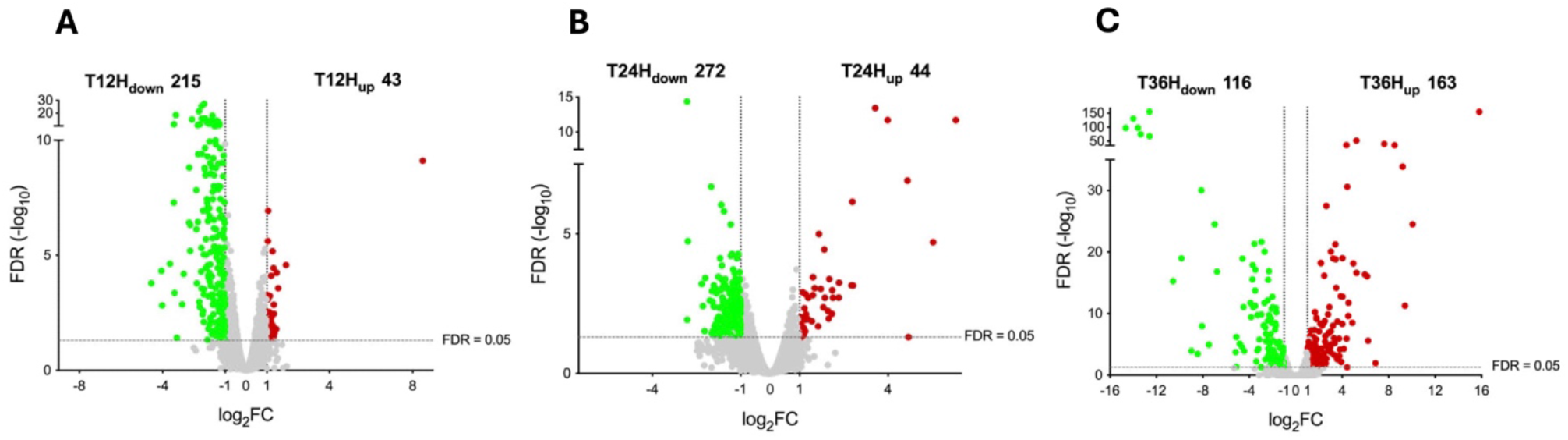
Differential gene expression analysis in mosquitoes with high versus low infection intensity. (A–C) Volcano plots showing differentially expressed (DE) genes between mosquitoes with high (H) and low (L) infection intensity at 12 (T12), 24 (T24), and 36 (T36) hpi. The x-axis represents log₂ fold change (log₂FC), and the y-axis shows −log₁₀FDR. Vertical dashed lines indicate log₂FC thresholds (-1 and 1), and the horizontal dashed line indicates the FDR significance cutoff (0.05). Upregulated genes (H vs. L) are shown in red, downregulated genes in green, and genes with non-significant changes in grey. The number of significantly up- and downregulated genes is indicated in each panel.

The complete lists of upregulated and downregulated genes for each comparison (12H vs 12L, 24H vs 24L, and 36H vs 36L), together with their functional annotations, are provided in the Supporting Excel File S3.

Gene Ontology (GO) enrichment analysis of the 215 genes downregulated in the H sample highlighted that 12 hours post-infection (hpi) represents the time point most responsive to differences in parasite burden in terms of immune activation (Figure S9). In particular, mosquitoes with low *Plasmodium* infection intensity exhibited strong and significant enrichment of GO terms associated with immune-related biological processes (e.g., regulation of innate immune responses, antimicrobial peptide production, activation of the melanization cascade) providing independent functional evidence for an early, robust, and coordinated immune response in mosquitoes carrying lower parasite loads. At 24 hours post-infection, GO enrichment analysis of the 272 genes downregulated in the H sample revealed a different transcriptional landscape in mosquitoes with low infection intensity, characterized by significant enrichment of terms related to cell–cell communication and signal transduction (Figure S9). This pattern suggests a transition from an early immune-effector phase toward broader regulatory and signalling processes as the infection progresses and parasite invasion of the midgut epithelium continues. Finally, at 36 hours post-infection, GO terms significantly enriched in mosquitoes with high parasite loads, corresponding to the onset of early oocyst development, were primarily associated with changes in fundamental physiological and metabolic processes. This indicates that, at this stage, the mosquito transcriptional response is less centered on immune defence and more reflective of systemic physiological adjustments to sustained infection (Supporting File S1, Figure S9).

### Stronger mosquito immune activation correlates with lower *Plasmodium* infection intensity

Analysis of the DE gene lists combined with GO enrichment analysis revealed significant modulation of immune-related genes and gene families at the early time points (12 and 24 hpi). Mosquitoes harbouring different parasite burdens exhibited markedly different levels of immune- related transcripts, with the greatest divergence observed at 12 hpi. Among the 215 genes downregulated in mosquitoes with high infection intensity at 12 hpi (i.e., upregulated in low- infection mosquitoes), 87 genes (40.5%) belonged to immune-related families. In contrast, only 9 of the 43 genes (20.1%) upregulated in high-infection mosquitoes were associated with immune functions (Supporting Excel File S3). This striking asymmetry indicates a substantially stronger and broader immune activation and/or transcriptional persistency in mosquitoes carrying lower parasite loads. The most parsimonious interpretation of our results is that a subset of mosquitoes mounts a rapid and effective anti-*Plasmodium* immune response, thereby limiting parasite development. Conversely, other mosquitoes, possibly owing to intrinsic physiological differences, fail to activate a sufficiently robust response, resulting in higher infection intensities. Consistent with this interpretation, among the genes upregulated in mosquitoes with low infection intensity at 12 hpi, we identified most of the genes previously described as key components of the *Anopheles* immune response against *Plasmodium* (Table 5).

**Table 5.** Heat map reporting the expression profiles in the different samples of the 87 immune-related genes upregulated at 12 hpi in the T12_L sample.

| ID | log<sub>2</sub>FC | FDR | Gene Name | Product Description/Pfam Description | T12_H | T12_L | T24_H | T24_L | T36_H | T36_L |
|---|---|---|---|---|---|---|---|---|---|---|
| **AGAP008783** | **-2.140922943** | **2.86E-26** | **N/A** | **Arginase** |  |  |  |  |  |  |
| AGAP008141 | -1.674696339 | 0.00724591 | N/A | argininosuccinate lyase |  |  |  |  |  |  |
| AGAP003015 | -2.095543695 | 2.86E-16 | N/A | Argininosuccinate synthase |  |  |  |  |  |  |
| AGAP013120 | -1.565413297 | 0.000888952 | Cht5-3 | chitinase |  |  |  |  |  |  |
| AGAP013166 | -1.008842481 | 0.001216551 | Cht5-1 | chitinase |  |  |  |  |  |  |
| AGAP009033 | -1.407014378 | 7.76E-15 | HPX2 | heme peroxidase 2 |  |  |  |  |  |  |
| AGAP005335 | -1.183631388 | 0.000180215 | CTL4 | C-type lectin (CTL) |  |  |  |  |  |  |
| AGAP005334 | -1.191435418 | 5.40E-05 | CTLMA2 | C-type lectin (CTL) - mannose binding |  |  |  |  |  |  |
| AGAP007407 | -1.418766724 | 1.21E-05 | CTLMA4 | C-type lectin (CTL) - mannose binding |  |  |  |  |  |  |
| AGAP004847 | -1.305713499 | 0.001249746 | SCRB7 | Class B Scavenger Receptor (CD36 domain). |  |  |  |  |  |  |
| AGAP005625 | -1.101785003 | 0.000583539 | SCRASP1 | Class A Scavenger Receptor (SRCR domain), Serine Protease |  |  |  |  |  |  |
| **AGAP001826** | **-1.173049905** | **0.009693342** | **Lp** | **lipophorin** |  |  |  |  |  |  |
| AGAP008193 | -1.109806401 | 3.68E-09 | N/A | Nidogen (entactin) |  |  |  |  |  |  |
| AGAP029385 | -1.401779013 | 8.65E-13 | N/A | Lipocln_cytosolic_FA-bd_dom domain-containing protein |  |  |  |  |  |  |
| AGAP002593 | -1.269623552 | 3.85E-12 | N/A | outer membrane lipoprotein Blc |  |  |  |  |  |  |
| AGAP005072 | -1.408164154 | 1.61E-06 | N/A | Laminin G domain |  |  |  |  |  |  |
| AGAP005848 | -2.009554902 | 4.94E-28 | N/A | ficolin A |  |  |  |  |  |  |
| AGAP009556 | -1.529387995 | 3.70E-11 | N/A | Fibrinogen domain |  |  |  |  |  |  |
| AGAP000806 | -1.289123629 | 4.64E-06 | N/A | Angiopoietin-like 1 |  |  |  |  |  |  |
| AGAP009107 | -1.721418482 | 1.00E-09 | N/A | Beta-1,3-glucan-binding protein, N-terminal |  |  |  |  |  |  |
| AGAP009109 | -1.167073675 | 0.003128032 | N/A | Beta-1,3-glucan-binding protein, N-terminal |  |  |  |  |  |  |
| **AGAP006275** | **-2.305040824** | **4.01177E-10** | **N/A** | **27 kDa Hemolymph Proteins** |  |  |  |  |  |  |
| AGAP007033 | -1.713685252 | 2.87E-09 | APL1C | Anopheles Plasmodium-responsive Leucine-Rich Repeat 1C |  |  |  |  |  |  |
| AGAP006348 | -2.063518142 | 1.15E-07 | LRIM1 | leucine-rich immune protein (Long) |  |  |  |  |  |  |
| AGAP007037 | -1.050778892 | 7.41E-07 | LRIM3 | leucine-rich immune protein (Long) |  |  |  |  |  |  |
| AGAP007039 | -1.969968925 | 1.11E-16 | LRIM4 | leucine-rich immune protein (Long) |  |  |  |  |  |  |
| AGAP027997 | -1.443702529 | 3.16E-11 | LRIM5 | leucine-rich immune protein (Short) |  |  |  |  |  |  |
| AGAP007454 | -1.346564391 | 5.29E-06 | LRIM8A | leucine-rich immune protein (Short) |  |  |  |  |  |  |
| AGAP007456 | -1.536593429 | 6.75E-07 | LRIM8B | leucine-rich immune protein (Short) |  |  |  |  |  |  |
| AGAP007453 | -1.176355406 | 0.001813429 | LRIM9 | leucine-rich immune protein (Short) |  |  |  |  |  |  |
| AGAP007455 | -1.238081463 | 6.29E-05 | LRIM10 | leucine-rich immune protein (Short) |  |  |  |  |  |  |
| AGAP007045 | -2.034149483 | 9.17E-06 | LRIM15 | leucine-rich immune protein (TM) |  |  |  |  |  |  |
| **AGAP005693** | **-1.526493557** | **3.36E-09** | **LRIM17** | **leucine-rich immune protein (Coil-less)** |  |  |  |  |  |  |
| AGAP002542 | -2.082891458 | 0.00367358 | LRIM20 | leucine-rich immune protein (Coil-less) |  |  |  |  |  |  |
| AGAP013290 | -1.603201757 | 1.71E-08 | LRIM27 | leucine-rich immune protein (Coil-less) |  |  |  |  |  |  |
| AGAP003878 | -1.173996511 | 4.43E-05 | N/A | Leucine-rich repeat |  |  |  |  |  |  |
| AGAP007457 | -2.989565191 | 6.46E-05 | N/A | Leucine-rich repeat |  |  |  |  |  |  |
| AGAP013186 | -1.280472985 | 0.000493923 | N/A | Leucine-rich repeat |  |  |  |  |  |  |
| AGAP004832 | -1.635909075 | 0.026844929 | N/A | Leucine-rich repeat;Leucine rich repeat 4;Leucine-rich repeat |  |  |  |  |  |  |
| AGAP011791 | -2.318416832 | 3.54E-07 | CLIPA1 | CLIP-domain serine protease |  |  |  |  |  |  |
| AGAP011788 | -1.101247862 | 1.00E-09 | CLIPA14 | CLIP-domain serine protease |  |  |  |  |  |  |
| AGAP011790 | -1.835038548 | 3.29E-07 | CLIPA2 | CLIP-domain serine protease |  |  |  |  |  |  |
| AGAP001964 | -1.152309265 | 0.000214724 | CLIPA26 | CLIP-domain serine protease |  |  |  |  |  |  |
| AGAP000290 | -1.157909877 | 2.00E-05 | CLIPA27 | CLIP-domain serine protease |  |  |  |  |  |  |
| AGAP011780 | -1.236991123 | 2.26E-10 | CLIPA4 | CLIP-domain serine protease |  |  |  |  |  |  |
| **AGAP011792** | **-1.920618025** | **4.36E-15** | **CLIPA7** | **CLIP-domain serine protease** |  |  |  |  |  |  |
| AGAP010731 | -1.99320389 | 1.62E-09 | CLIPA8 | CLIP-domain serine protease |  |  |  |  |  |  |
| **AGAP010968** | **-1.840958856** | **2.73E-05** | **CLIPA9** | **CLIP-domain serine protease** |  |  |  |  |  |  |
| AGAP003251 | -1.281904332 | 2.48E-11 | CLIPB1 | CLIP-domain serine protease |  |  |  |  |  |  |
| **AGAP009844** | **-1.420957795** | **7.40E-06** | **CLIPB15** | **CLIP-domain serine protease** |  |  |  |  |  |  |
| AGAP003247 | -1.36699358 | 3.60E-05 | CLIPB19 | CLIP-domain serine protease |  |  |  |  |  |  |
| AGAP012037 | -1.797427978 | 9.52E-05 | CLIPB20 | CLIP-domain serine protease |  |  |  |  |  |  |
| AGAP013184 | -1.030009639 | 0.006865118 | CLIPB36 | CLIP-domain serine protease |  |  |  |  |  |  |
| AGAP003250 | -1.397385498 | 2.11E-11 | CLIPB4 | CLIP-domain serine protease |  |  |  |  |  |  |
| **AGAP003057** | **-1.639145756** | **8.19E-05** | **CLIPB8** | **CLIP-domain serine protease** |  |  |  |  |  |  |
| AGAP004719 | -1.342241315 | 0.000176649 | CLIPC9 | CLIP-domain serine protease |  |  |  |  |  |  |
| **AGAP028725** | **-1.933168111** | **3.54E-05** | **SPCLIP1** | **CLIP-domain serine protease** |  |  |  |  |  |  |
| AGAP028167 | -2.142012186 | 4.78E-06 | N/A | CLIP domain-containing serine protease |  |  |  |  |  |  |
| AGAP012034 | -1.589476603 | 0.001407911 | N/A | CLIP domain-containing serine protease |  |  |  |  |  |  |
| AGAP012900 | -1.165874675 | 1.25E-06 | N/A | Peptidase C1A, papain C-terminal |  |  |  |  |  |  |
| AGAP002720 | -1.948657398 | 3.36E-09 | N/A | cathepsin O |  |  |  |  |  |  |
| AGAP003926 | -1.210411413 | 0.00051296 | N/A | Aminopeptidase |  |  |  |  |  |  |
| AGAP011475 | -1.22152325 | 3.33E-05 | N/A | Envelysin |  |  |  |  |  |  |
| **AGAP008371** | **-1.807082645** | **0.015725415** | **N/A** | **Peptidase_M14 domain-containing protein** |  |  |  |  |  |  |
| AGAP007043 | -2.130965226 | 1.72E-16 | N/A | Serine proteases, trypsin domain |  |  |  |  |  |  |
| AGAP003248 | -1.864688901 | 5.21E-08 | N/A | Serine proteases, trypsin domain |  |  |  |  |  |  |
| **AGAP013221** | **-1.176252553** | **1.82E-07** | **N/A** | **Serine proteases, trypsin domain** |  |  |  |  |  |  |
| AGAP013252 | -2.140184913 | 0.000276644 | N/A | Serine proteases, trypsin domain |  |  |  |  |  |  |
| AGAP000182 | -1.233631155 | 0.006263991 | N/A | Serine proteases, trypsin domain |  |  |  |  |  |  |
| AGAP013117 | -1.218307574 | 0.013402787 | N/A | Serine proteases, trypsin domain |  |  |  |  |  |  |
| AGAP029126 | -1.153590147 | 0.013402787 | N/A | Serine proteases, trypsin domain |  |  |  |  |  |  |
| AGAP010545 | -1.020255956 | 0.014051605 | N/A | Serine proteases, trypsin domain |  |  |  |  |  |  |
| AGAP010730 | -1.853203682 | 5.06E-10 | N/A | prophenoloxidase activating factor |  |  |  |  |  |  |
| AGAP003245 | -3.061105367 | 0.001361249 | N/A | Serine proteases, trypsin domain;Proteinase, CLIP domain |  |  |  |  |  |  |
| **AGAP002268** | **-1.465540485** | **0.000268051** | **N/A** | **Thrombospondin type-1 (TSP1) repeat** |  |  |  |  |  |  |
| AGAP006910 | -1.031506174 | 5.06E-08 | SRPN3 | serine protease inhibitor (serpin) 3 |  |  |  |  |  |  |
| AGAP005620 | -1.217976584 | 0.000415977 | N/A | Attacin_C domain-containing protein |  |  |  |  |  |  |
| AGAP007385 | -1.245447089 | 0.000878435 | LYSC4 | C-type lysozyme |  |  |  |  |  |  |
| AGAP000274 | -1.168243349 | 0.000450345 | N/A | Invertebrate-type lysozyme |  |  |  |  |  |  |
| AGAP010815 | -1.696935456 | 0.000214724 | TEP1 | Thioester-containing protein 1 |  |  |  |  |  |  |
| AGAP010816 | -1.449805543 | 0.031301799 | TEP3 | thioester-containing protein 3 |  |  |  |  |  |  |
| AGAP010812 | -1.463559314 | 0.008240931 | TEP4 | thioester-containing protein 4 |  |  |  |  |  |  |
| AGAP010814 | -1.814602354 | 0.000596864 | TEP6 | thioester-containing protein 6 |  |  |  |  |  |  |
| AGAP008654 | -2.158476039 | 5.11E-05 | TEP12 | thioester-containing protein 12 |  |  |  |  |  |  |
| **AGAP008368** | **-1.505519573** | **2.48E-11** | **TEP14** | **thioester-containing protein 14** |  |  |  |  |  |  |
| AGAP010658 | -2.36250806 | 0.000162056 | N/A | Hemocyanin |  |  |  |  |  |  |
| AGAP012616 | -1.221587489 | 0.040711461 | PPO5 | prophenoloxidase 5 |  |  |  |  |  |  |

In bold, genes upregulated both at 12 and 24 hpi in mosquitoes harbouring low parasite load. The heat map reports the intensity of expression (TPM) in the different samples as indicated. Colour graduation indicates the level of expression: dark blue, higher TPM values, light blue or white, lower TPM values as indicated in the scale. Gene identification, log<sub>2</sub>FC, FDR, gene name (when applicable) and descriptions are reported.

Most immune genes upregulated in low-infection mosquitoes at 12 hpi are involved in parasite recognition and melanization response, a multi-step immune process classically organized into three phases: pathogen recognition, signal amplification and modulation, and effector activity. Among transcripts more abundant in low-infection mosquitoes, we identified 17 LRR- containing genes (including LRIM1, LRIM17, LRIM4, LRIM9, and LRIM8A) and six TEP family members (TEP1, TEP3, TEP4, TEP6, TEP12, and TEP14). LRIM proteins form complexes with TEPs and are essential for stabilizing complement-like immune responses against *Plasmodium*. Among genes involved in melanization, we observed strong enrichment of CLIP-domain serine proteases (CLIPs), together with their regulatory serine protease homologs (SPHs), C-type lectins (CTLs), and serine protease inhibitors (SRPNs). The CLIP family represents a central regulatory hub of the melanization cascade. In addition, we detected Class A and B scavenger receptors (SCRASP1 and SCRB7) and lipocalins implicated in immune priming. Finally, several downstream components of the melanization effector machinery were upregulated, including PPO5, a prophenoloxidase- activating factor, and multiple antimicrobial effectors such as C-type lysozymes. Together, these data indicate that mosquitoes carrying lower *Plasmodium* burdens mounted a markedly stronger, broader, and more coordinated immune response during the critical early phases of infection, strongly supporting the widely accepted idea that early immune activation plays a key role in limiting parasite establishment in the mosquito midgut. Furthermore, several components of the arginine metabolism and nitric oxide (NO) production pathway were transcriptionally upregulated in mosquitoes with low *Plasmodium* infection intensity. These include arginase (ARG), argininosuccinate lyase (ASL), and argininosuccinate synthase (ASS) (Table 5). In addition to arginine metabolism, several enzymes involved in uric acid metabolism, such as urate oxidase (AGAP008440), showed higher expression in mosquitoes with low infection intensity. These pathways are functionally linked to the production of nitric oxide (NO) and reactive oxygen species (ROS), both of which have been implicated in antiparasitic defense (Kwon and Smith, 2022).

Comparative analysis of differentially expressed (DE) gene sets across time points revealed a subset of genes (22 contigs, Figure 7B and Table 5) activated in mosquitoes with low infection intensity at both 12 and 24 hpi. Notably, 14 of the 22 annotated contigs (63.6%) shared between the DE gene sets upregulated in low-infection mosquitoes at 12 and 24 hpi were immune-related (Figure 7C).

**Figure 7.**
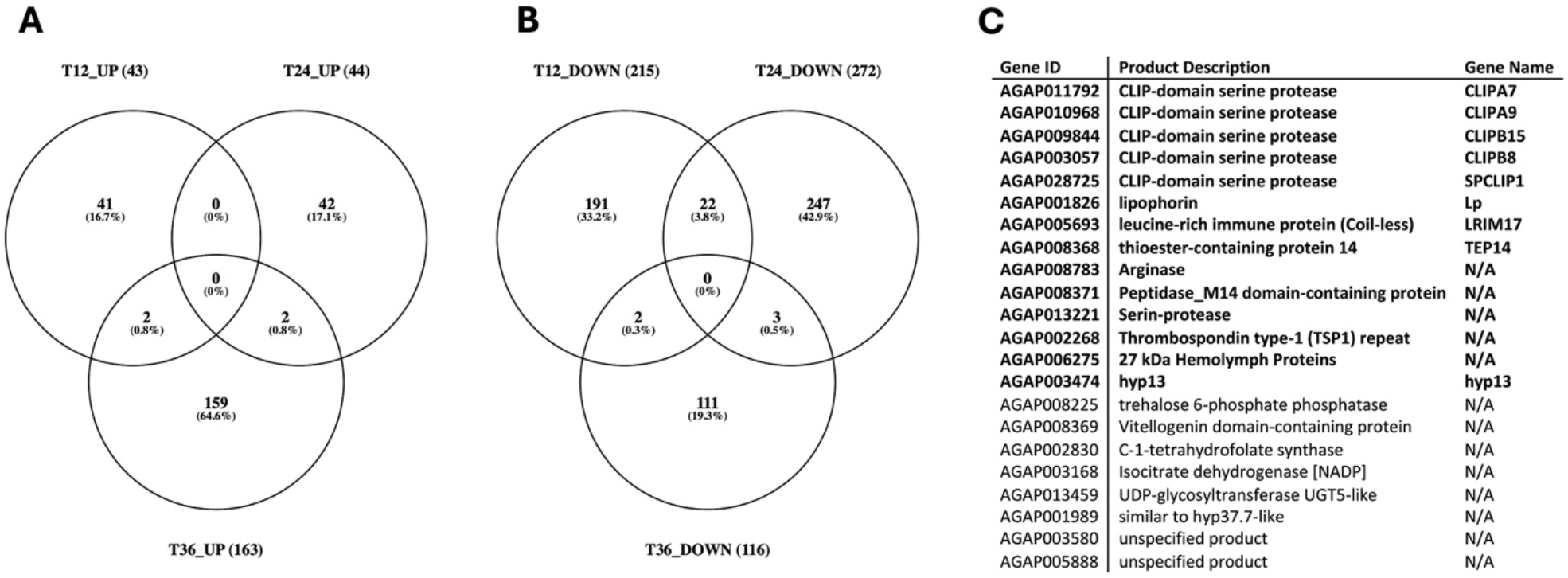
Overlap of differentially expressed mosquito genes across time points following infection. (A) Venn diagram showing overlap among genes upregulated in mosquitoes with high infection intensity at 12 h (T12_UP, 43 genes), 24 h (T24_UP, 44 genes), and 36 h (T36_UP, 163 genes) post-infection. (B) Venn diagram showing overlap among genes downregulated in high-infection mosquitoes at 12 h (T12_DOWN, 215 genes), 24 h (T24_DOWN, 272 genes), and 36 h (T36_DOWN, 116 genes) post-infection. (C) Table listing the group of 22 genes shared among downregulated gene sets at early time points (T12_DOWN and T24_DOWN); immune-related factors and genes involved in metabolic pathways relevant to immune defense, such as arginase, are in bold.

These observations clearly point to a sustained and progressive immune response spanning both early events, such as parasite escape from the peritrophic matrix, adhesion to the midgut epithelium and initial invasion, and later stages, involving epithelial traversal and early oocyst formation. Together, these results support the existence of sustained immune activation during the early and mid-phases of midgut invasion in mosquitoes with low *Plasmodium* infection intensity, which largely diminishes by 36 hpi, when most ookinetes have completed traversal of the midgut epithelium. In contrast, only a limited number of contigs were shared among the sets of genes upregulated in mosquitoes with high parasite loads across different time points, as shown in the Venn diagram (Figure 7A). This limited overlap suggests that transcriptional responses in high-infection mosquitoes are more transient, less coordinated, or more tightly restricted to specific stages of parasite development, rather than reflecting a sustained and coherent immune program (Figure 7 and Supporting Excel File S3). Moreover, transcriptional differences between groups were less pronounced at 36 hpi, suggesting that divergence in immune responses is most prominent during the early stages of mosquito–parasite interaction. Overall, these findings indicate that mosquitoes with lower parasite burdens not only activate classical immune pathways, including parasite recognition and melanization, but also upregulate metabolic pathways that may support the production of effector molecules such as NO and ROS, further reinforcing the association between metabolic reprogramming and effective antiparasitic immunity. Interestingly, other immune-related genes, such as the antimicrobial peptide hyp13 and the lipophorin were among the genes upregulated in mosquitoes with low infection intensity at both 12 and 24 hpi.

### Systemic modulation of mosquito immune gene families during *Plasmodium* development

To obtain possible additional information on variation of mosquito immune responses according to parasite burden, we extended the comparison between mosquitoes with low and high infection intensity to the entire transcriptome, rather than restricting it to differentially expressed (DE) genes. To identify immune-related transcripts and compare their expression levels (TPM) in our dataset, we leveraged the curated catalogue of *An. gambiae* immune genes previously described along with associated Gene Ontology (GO) and Pfam annotations (Neafsey et al., 2015). This catalogue includes 28 immune-related gene families involved in pathogen recognition, signaling pathways, cascade regulation, and effector mechanisms. First, using ImmunoDB (IPRO) IDs and annotations, we queried the *An. gambiae* AgamP4.14 genome (VectorBase release 57) and retrieved the AGAP identifiers of 922 genes belonging to immune- related families. Corresponding contigs and their TPM values were then extracted from our transcriptome dataset, retaining for downstream analysis only contigs with TPM > 1 in at least one sample (377 contigs, Supporting Excel File S4). To evaluate whether specific immune gene families or pathways were differentially associated with infection intensity, we calculated the proportion of transcripts within each family showing at least a 1.5-fold difference in expression between low- and high-intensity samples (Figure 8A–C). Several immune families displayed substantial transcriptional modulation, indicating that parasite burden is associated with changes across multiple components of the mosquito immune system. These differences were most pronounced at 12 and 24 hpi, whereas by 36 hpi the transcriptional profiles of immune genes between high- and low-intensity groups became less distinct.

**Figure 8.**
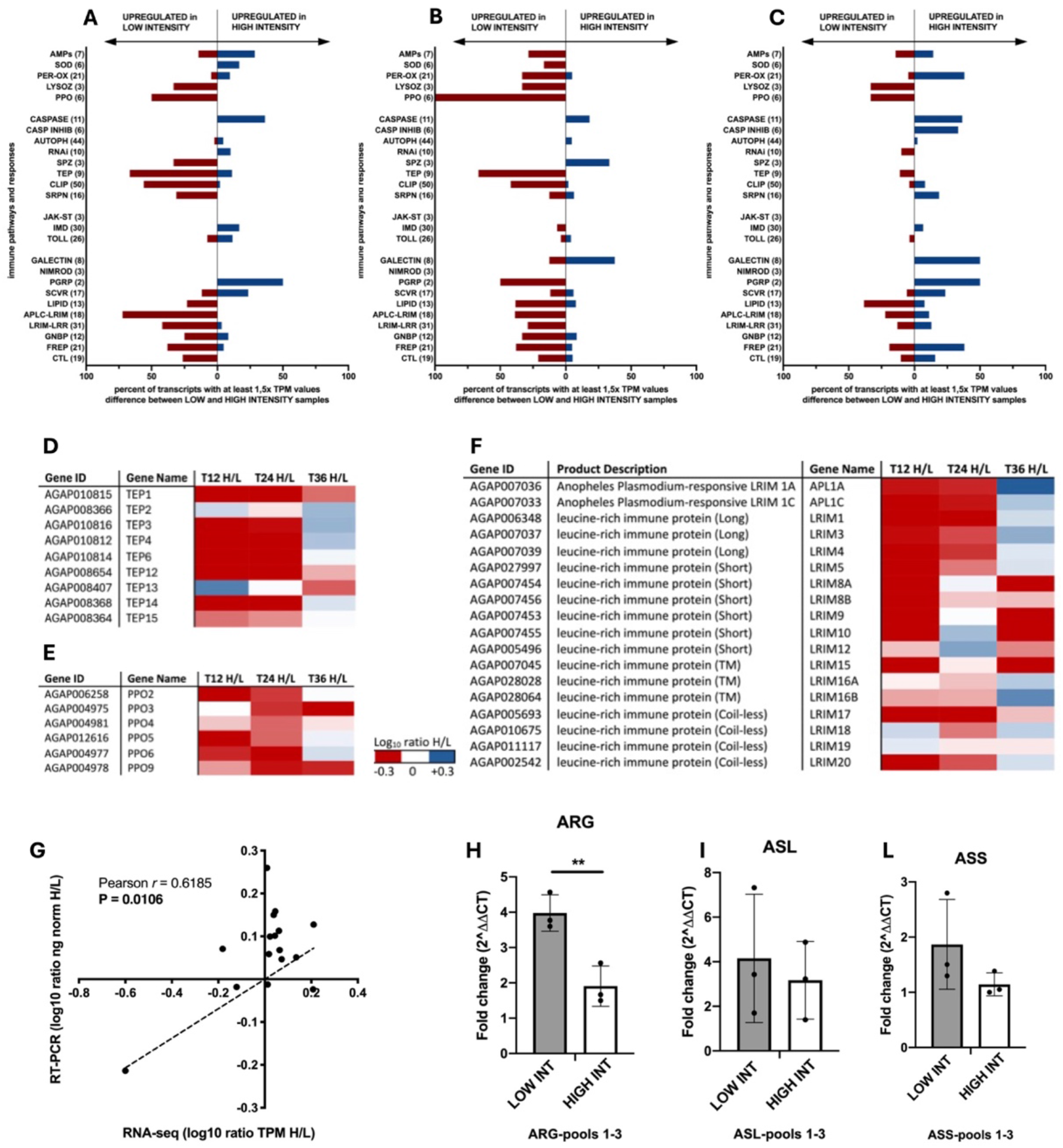
Differential modulation of mosquito immune pathways and validation by qPCR. (A–C) Relative abundance of immune-related transcripts in mosquitoes with low and high *P. falciparum* infection intensity at 12 (A), 24 (B), and 36 (C) hours post-infection (hpi). Bars represent the percentage of genes within each immune pathway or group showing ≥1.5-fold difference in TPM values. Immune pathways/groups are listed on the left y axis, with the total number of genes reported in brackets; left and right directions indicate genes upregulated in low- and high-intensity mosquitoes, respectively. (D–F) Heatmaps showing expression profiles (log₁₀ H/L TPM ratio) of selected immune gene families across time points: thioester-containing proteins (TEP) (D), prophenoloxidases (PPO) (E), and leucine-rich immune proteins (LRIM) (F). (G) Correlation between RNA-seq and qPCR expression measurements for the genes selected for qPCR analysis: SCRBQ2, ANXB9, LRR, Peritrophin-A, Carboxypeptidase B and Ficolin-A. Dots represent averaged RNA-seq and qPCR values as indicated in the x- and y-axes for the five genes at 12, 24 and 36 hpi. (H–L) qPCR validation of genes involved in arginine metabolism, i.e., arginase (ARG) (H), argininosuccinate lyase (ASL) (I), and argininosuccinate synthase (ASS) (L), in pools of mosquitoes with low and high infection intensity at 12 hpi. Data are shown as fold change (2^-ΔΔCT); bars represent mean ± SD. **: P < 0.01.

Overall, observations from this extended analysis are consistent with the patterns identified by differential expression (DE) analysis. Gene families involved in pathogen recognition exhibited particularly strong transcriptional differences between infection intensities. These included families encoding peptidoglycan recognition proteins (PGRPs), C-type lectins (CTLs), fibrinogen-related proteins (FREPs), and leucine-rich repeat immune proteins (LRIMs). These proteins act as pattern-recognition receptors, detecting microbial molecules and parasite surfaces and initiating downstream immune responses. In contrast, core immune signaling pathways, including components of the Toll, IMD, and JAK/STAT pathways, showed relatively limited transcriptional variation between infection groups. This is consistent with previous observations that many signaling components are constitutively expressed and primarily regulated through upstream activation or post-transcriptional mechanisms (Cirimotich et al., 2010; Clayton et al., 2014; Simões et al., 2017). Genes involved in protease cascade regulation also displayed transcriptional modulation at 12 and 24 hpi, including CLIP-domain serine proteases, serine protease inhibitors (SRPNs), and Spaetzle (SPZ) proteins, which contribute to melanization and immune signal amplification. Several key immune effectors were differentially expressed between infection intensities. Several members of the thioester-containing protein (TEP) family showed higher transcriptional abundance in mosquitoes with low-infection intensity (Figure 8D). TEPs function as complement-like factors and play a central role in parasite killing. Similarly, genes encoding prophenoloxidases (PPOs), key components of the melanization pathway, exhibited consistent transcriptional patterns across all three time points, with higher expression in low- infection mosquitoes (Figure 8E). A particularly strong response was observed in the LRIM and APL1 gene families (Figure 8F) in mosquitoes with low-infection intensity, as also shown in DE analysis. In contrast, gene families associated with metabolic stress and cellular damage, including autophagy-, apoptosis- (caspases), and oxidative stress–related genes (peroxidases), showed higher transcriptional abundance in mosquitoes with high infection intensity, across the three time points.

To validate the RNA-seq results, expression profiles of a subset of genes was analyzed by real time qPCR, specifically Class B Scavenger Receptor (SCRBQ2, Croquemort Homologue, AGAP010133), Annexin B9 (ANXB9, AGAP003790), Leucine-Rich Repeat (LRR, AGAP006644), Chitin Binding Peritrophin A (AgAper25a, AGAP010364), Carboxypeptidase B (cpbAg1, AGAP006209), Ficolin-A (AGAP005848), Argininosuccinate synthase (ASS, AGAP003015), Argininosuccinate lyase (ASL, AGAP008141) and Arginase (ARG, AGAP008783). Expression levels obtained by the two methods were significantly correlated (Pearson r = 0.6185, P = 0.0106), confirming the robustness of the transcriptomic analysis (Figure 8G). Also, qPCR assays further confirmed differential expression of genes involved in arginine metabolism between infection intensity groups (Figure 8H–L).

## DISCUSSION

In this study, we established a molecular framework to quantify *P. falciparum* ookinetes in individual *An. coluzzii* mosquitoes and to associate parasite burden with mosquito immune responses. By focusing on the ookinete stage, a critical developmental bottleneck in the parasite life cycle, we aimed to provide new insights into the early mosquito–parasite interactions that are likely to influence infection outcome and transmission potential. Time-resolved transcriptomic analysis (12–36 hours post-infectious blood meal) revealed distinct mosquito transcriptional states associated with low and high parasite burdens during ookinete maturation and midgut invasion.

First, we validated four microneme-associated genes (ctrp, warp, soap, and cht1) as reliable molecular markers for detecting and quantifying ookinetes in individual mosquitoes. These genes, which are involved in motility, epithelial adhesion, and midgut invasion, showed stage- specific expression patterns consistent with previous studies (Dessens et al., 1999, 2003; Langer et al., 2000; Abraham et al., 2004; Vlachou et al., 2004, 2006; Akinosoglou et al., 2015; Real et al., 2021). We used laboratory-reared *An. coluzzii* mosquitoes infected with a laboratory strain of *P. falciparum*, with the infectious blood meal carefully controlled to minimize confounding factors such as mosquito age, microbiota composition, blood meal variability, and gametocyte density. qPCR approach confirmed peak expression of *Plasmodium* markers between 18 and 24 hpi.

Absolute quantification indicated that all four markers could serve as proxies for ookinete abundance, although soap and cht1 consistently exhibited higher transcript levels. These differences may reflect variation in gene expression, transcript stability, or assay sensitivity. Compared to traditional infectivity assays based on oocyst or sporozoite detection, which require several days, our approach enables estimation of parasite burden within 12–36 hours post- infection and allows the analysis of mosquito nucleic acids in the same samples used for ookinete quantification. The robust concordance observed between ookinete transcript abundance and subsequent oocyst counts in the infection experiment further supports the use of these markers as early molecular proxies of infection intensity. A first key observation of this study is the substantial inter-individual variability in parasite load among mosquitoes exposed to identical gametocytemia under controlled conditions. The consistency of parasite load estimates in individual mosquitoes was supported by strong correlations among the four ookinete markers, enabling stratification of mosquitoes into low- and high-burden groups.

To date, most studies on mosquito innate immunity have focused on comparisons between infected and uninfected individuals, or between mosquitoes exposed to high versus low gametocytemia, to identify genes activated by the parasite and involved in antiparasitic responses (Cohuet et al., 2006; Mendes et al., 2011; Mead et al., 2012; Simões et al., 2017). Moreover, most studies investigating the immune response of *An. gambiae* to malaria parasites have relied on the rodent malaria model *Plasmodium berghei* (Dong et al., 2006; Severo and Levashina, 2014). In contrast, we employed transcriptomic analysis to investigate how mosquito immune responses to *P. falciparum* vary over the course of ookinete development and in relation to parasite infection intensity. A novelty of our study is the stratification of mosquitoes based on accurately quantified ookinete loads, enabling direct comparison between individuals with high and low *Plasmodium* infection intensities. We have employed the mosquitoes singularly characterized to quantify the *P. falciparum* ookinete-associated transcripts to investigate within-infection variability in mosquito immune responses. Differential expression and gene ontology enrichment analyses allowed to identify genes associated with early *Anopheles*–*P. falciparum* interactions and to correlate alterations of immune-related pathways with infection intensity. The analysis of *Plasmodium* transcriptional profiles at different times post-infection revealed that relatively few parasite genes were differentially expressed between conditions, indicating limited transcriptional plasticity in response to infection intensity at these early stages. Among parasite genes upregulated at high infection intensity at 12 hpi were markers associated with motility, invasion, structural integrity, and immune evasion, including IMC proteins, Cap380, G377, SIMP, α-tubulin, PRP8, and PIMMS43. Conversely, downregulated genes included components of chromatin organization and ribosomal machinery. These changes may reflect resource reallocation toward invasion and differentiation processes under conditions of increased parasite density (Supporting Excel File S5).

On the contrary, mosquito transcriptional responses differed markedly between infection groups. Mosquitoes with lower parasite burdens exhibited stronger and more consistent transcriptional responses at 12 and 24 hpi, corresponding to the early stages of P. falciparum development within the mosquito. Notably, 14 of the 22 annotated contigs (63.6%) shared between the sets of genes upregulated in low-infection mosquitoes at both time points were immune-related, suggesting a prominent and sustained involvement of immune pathways during early parasite development. This response involved genes associated with parasite recognition (e.g., LRRs, CTLs, scavenger receptors), complement-like system activation (TEP family and LRIM/APL1 complex), and melanization (CLIP proteases and PPOs), all of which are well- established components of anti-*Plasmodium* immunity (Christophides et al., 2002; Osta et al., 2004a; Yassine and Osta, 2010; Volohonsky et al., 2020). In *An. gambiae*, parasite recognition is largely mediated by leucine-rich repeat (LRR) proteins and complement-like thioester-containing proteins (TEPs). In particular, LRIM1 and APL1C are known to form a complex with TEP1 (the TEP1/LRIM1/APL1C complex), which binds invading ookinetes and mediates their elimination, either by direct lysis or by triggering melanization (Blandin et al., 2004; Fraiture et al., 2009; Povelones et al., 2009, 2011). We also found several members of the serine protease cascade upregulated in low-infection mosquitoes, including CLIPB1, CLIPB4, CLIPB8, and CLIPB15 (Zhang et al., 2023), as well as negative regulators (CLIPA2, CLIPA5, and CLIPA7) and positive regulators such as CLIPC9 and CLIPA8, which are essential for activating melanization downstream of TEP1 binding (Sousa et al., 2020). More recently, SPCLIP1 has been shown to be required for stable deposition of full-length TEP1 onto *Plasmodium* ookinetes, acting downstream of initial parasite recognition. We also identified three C-type lectins: CTL4 and CTLMA2, which are known agonists of parasite development because they negatively regulate melanization (Osta et al., 2004b, 2004a; Simões et al., 2022), and CTLMA4. The enrichment of these pathways at early time points suggests that lower parasite burdens are associated with the rapid and sustained activation of multiple immune mechanisms during ookinete development, attachment to the midgut epithelium, and subsequent invasion. A significant inverse correlation between developing oocysts and melanized ookinetes has been previously reported (Lombardo and Christophides, 2016), suggesting that enhanced melanization may limit parasite development, whereas reduced melanization could be associated with higher parasite burden. Other interesting immune-related factors that were differentially regulated during the early stages of *Plasmodium* development (i.e., at both 12 and 24 hpi; Figure 7) include lipophorin (Cheon et al., 2006), the 27 kDa Hemolymph Protein (Muñoz-Gómez et al., 2014) and hyp13 (Arcà et al., 2017; Bevivino et al., 2021, 2024). Lipophorin and the 27 kDa hemolymph protein have both been implicated in insect immune responses, while hyp13 has only recently been reported to possess antimicrobial activity against bacteria (Bevivino et al., 2024).

Interestingly, previous transcriptomic studies also identified hyp13 as being upregulated following *Plasmodium berghei* infection (Vlachou et al., 2005).

In addition to canonical immune pathways, we observed differential modulation of genes involved in L-arginine and uric acid metabolism, including ARG, ASS, ASL, and urate oxidase. ARG catalyzes the conversion of L-arginine into L-ornithine and urea in the final step of the urea cycle. In contrast, ASS converts citrulline and aspartate into argininosuccinate, which is subsequently cleaved by ASL to generate fumarate and regenerate L-arginine. Together, these enzymes function to replenish intracellular L-arginine pools. This is particularly relevant for mosquito antiparasitic immunity, as L-arginine serves as the substrate for nitric oxide synthase (NOS), which produces nitric oxide (NO), a potent effector molecule with well-established anti- *Plasmodium* activity (Yuan et al., 2017). Notably, urate oxidase (AGAP008440), which catalyzes uricolysis, the conversion of uric acid into glyoxylic acid and urea, was significantly upregulated. This enzyme was previously shown to be induced following *Plasmodium* infection and to contribute to parasite killing, as its silencing leads to elevated uric acid levels and enhanced parasite survival (Kwon and Smith, 2022). We also observed upregulation of xanthine dehydrogenase (AGAP007918), the enzyme that catalyzes the oxidation of xanthine to uric acid. In *Drosophila*, this enzyme is active in the gut and has been implicated in regulating reactive oxygen species (ROS) and NO production, thereby exerting a protective immune function (Kim et al., 2001). The coordinated regulation of these metabolic genes suggests that metabolic reprogramming may accompany immune activation and contribute to parasite restriction, highlighting a potential interplay between metabolism and immunity in the mosquito response to infection.

The differential expression of immune-related genes observed as early as 12 hpi suggests that mosquito immune responses may be initiated during the early stages of *P. falciparum* development. At this time point, the observed response likely reflects the combined effects of developing ookinetes still present in the blood bolus and those beginning to interact with the midgut epithelium (Siciliano et al., 2020b) . Previous studies have reported increased expression of mosquito genes, including carboxypeptidase A and Defensin, as early as 14 hours post-blood meal, coinciding with the early stages of ookinete maturation within the midgut lumen (Tahar et al., 2002; Lavazec et al., 2007). Consistent with these findings, the expression patterns observed in the L and H groups between 12 and 24 hpi, particularly for immune-related genes, suggest that differences in mosquito immune responses may emerge early during parasite development, potentially before ookinete attachment to and traversal of the midgut epithelium. The higher expression of immune-related genes observed in mosquitoes with lower parasite loads may reflect an early protective response that limits parasite development or, alternatively, pre-existing differences in immune competence that influence susceptibility to infection. However, natural variation in mosquito responses to blood feeding may also contribute to the observed differences. Further functional studies will be required to distinguish between these possibilities and establish whether early immune activation plays a causal role in limiting parasite development. Temporal analysis further indicated that the pronounced immune-related transcriptional differences between low- and high-infection mosquitoes during early stages (12–24 hpi) became less distinct by 36 hpi, when parasites develops toward oocyst formation. This pattern is consistent with the idea that early immune responses are particularly critical in determining infection outcome, whereas later stages may reflect more generalized physiological adjustments.

Interestingly, mosquitoes with high parasite loads displayed increased abundance of genes associated with cellular stress, apoptosis, autophagy, and oxidative damage. This pattern may reflect either a reduced capacity to mount an effective immune response or a physiological consequence of increased parasite burden, potentially resulting from more extensive tissue damage during ookinete traversal of the midgut epithelium. Notably, because all mosquitoes were exposed to the same parasite challenge, these differences are more likely to reflect variability in mosquito responsiveness rather than differences in exposure. Previous studies have reported increased immune activation under high parasite exposure (Mendes et al., 2011; Garver et al., 2012; Simões et al., 2017). In contrast, our experimental design controlled for gametocytemia, allowing us to isolate variation in vector response. Our findings therefore support a model in which differences in infection outcome are also associated with variation in mosquito immune activation rather than only on differences in parasite input.

In summary, this study establishes a robust molecular approach for the early quantification of *P. falciparum* infection at the ookinete stage. An important aspect of our experimental design was the use of a single, standardized infection to investigate the relationship between ookinete burden and mosquito transcriptional responses at the individual level. By exposing mosquitoes to the same infectious blood meal under controlled experimental conditions, we minimized variation associated with parasite gametocytemia, blood composition, and infection procedures, allowing us to focus on differences in parasite development among individual mosquitoes. This approach was particularly relevant for investigating whether mosquitoes exposed to identical infection conditions but exhibiting different parasite burdens also displayed distinct transcriptional responses. Nevertheless, the use of a single infection experiment limits the generalizability of our findings, as infection-specific effects cannot be excluded. Further studies involving independent infections will therefore be necessary to establish the reproducibility of the observed transcriptional patterns across different experimental conditions.

The correlation between molecular quantification of ookinete-specific transcripts and microscopy-based oocyst counts provides a link between early parasite burden and subsequent oocyst development, a well-established indicator of mosquito infection intensity. Importantly, during the first 36 hpi, nucleic acids from the mosquito, parasite, and human blood meal coexist, offering a unique opportunity to investigate host–parasite–vector interactions within individual mosquitoes. In particular, the persistence of nucleic acids from human nucleated blood cells during this early window may enable the simultaneous characterization of human host genetic factors, parasite development, and mosquito responses. This integrated approach holds considerable potential for both laboratory investigations and field studies aimed at identifying determinants of malaria transmission.

Finally, we found that mosquito transcriptional responses were associated with parasite infection intensity. Mosquitoes with lower parasite burdens exhibited stronger early expression of immune- and metabolism-related genes, whereas higher parasite burdens were associated with weaker transcriptional responses. These findings suggest that early mosquito immune and metabolic responses may contribute to variation in susceptibility to P. falciparum infection and highlight the importance of early vector–parasite interactions in determining infection outcomes.

## SUPPORTING INFORMATIONS

### Supporting File S1: Figure S1-S10, Table S1

**Supporting Excel File S1: Composition of mosquito pools used for RNA-seq analysis.** Detailed description of mosquito samples included in each RNA pool used for transcriptomic analysis. For each pool (T12, T24, and T36; low (L) and high (H) infection intensity), the table reports the individual mosquito identifiers, corresponding ookinete marker expression values (ctrp, soap, warp, and cht1) quantified by qPCR (calculated using single-experiment standard curves).

Mosquitoes were assigned to low- or high-infection categories based on consistent transcript levels across at least three of the four markers. Each pool is composed by mosquitoes highlighted with the same color (ctrp and cht1 are the markers used for the "high" and "low" intensity classification).

**Supporting Excel File S2: Transcriptome with annotation and expression profiles.** List of *An. gambiae* genes (AGAP IDs) with corresponding functional annotations and expression levels in mosquitoes with high (H) and low (L) *P. falciparum* infection intensity at 12, 24, and 36 hours post- infection (hpi). Columns report gene identifier, product description, gene name or symbol (when available), Pfam domain IDs and descriptions, and transcript abundance values. Expression levels are shown as TPM values averaged for each condition (T12_H, T12_L, T24_H, T24_L, T36_H, T36_L). Ratios (H/L) indicate relative expression between high- and low-infection groups at each time point, and corresponding log-transformed values (log₂ H/L) are provided.

List of P. falciparum genes (PF3D7 IDs) with corresponding transcript abundance values. Expression levels are shown as TPM values averaged for each condition (T12_H, T12_L, T24_H, T24_L, T36_H, T36_L).

**Supporting Excel File S3: Differentially expressed (DE) *An. gambiae* genes.** Complete lists of differentially expressed (DE) genes identified by RNA-seq analysis in *An. gambiae* mosquitoes comparing high (H) versus low (L) *P. falciparum* infection intensity at 12, 24, and 36 hours post- infection (hpi). For each gene, the table reports AGAP gene ID, gene name and functional annotation, log₂ fold change (H/L) and statistical significance values (P-value and FDR).

**Supporting Excel File S4: Immune catalogue and immune gene expression values in the transcriptome.** The tables summarize the expression profiles of genes belonging to immune families in *An. gambiae* across different infection conditions and time points. Gene IDs and product descriptions are indicated, transcript abundance (TPM) is also reported for mosquitoes with high (H) and low (L) *P. falciparum* infection intensity at 12, 24, and 36 hours post-infection (hpi). For each gene, relative expression between conditions is represented as H/L ratios and corresponding log₂(H/L) values.

**Supporting Excel File S5: Differentially expressed (DE) *P. falciparum* genes.** Complete lists of *P. falciparum* differentially expressed (DE) genes identified by RNA-seq analysis in *An. gambiae* mosquitoes comparing high (H) versus low (L) P. falciparum infection intensity at 12, 24, and 36 hours post-infection (hpi). For each gene, the table reports PF3D7 gene ID, gene name and functional annotation, logarithmic transformation (log₂) of fold change (H/L) and CPM (counts per million reads) values and statistical significance values (P-value and FDR). Essential features of known ookinete factors are reported (highlighted in green). Markers used in this work are highlighted in yellow.

## ACKNOWLEDGEMENTS

We thank Paola Serini for her assistance with mosquito rearing. We are also grateful to Giulia Costa and Elena Levashina (Max Planck Institute for Infection Biology, Berlin, Germany) for providing *Anopheles coluzzii* strain N’Gousso mosquitoes infected with *Plasmodium falciparum* through the Infravec project (#5876).

