## Supplementary_Figures_S1 for "Molecular quantification of *Plasmodium falciparum* ookinete load reveals associations with early immune responses in *Anopheles coluzzii* mosquitoes"

#### Transcriptional profiling of ookinete markers in pools of infected mosquitoes

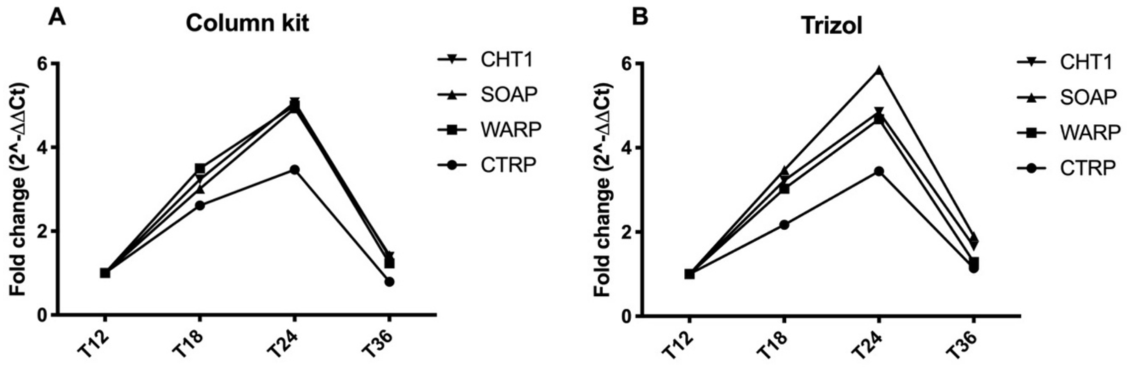

**Figure S1. Ookinete stage identification and markers transcriptional analysis.** The expression profile of *ctrp*, *warp*, *soap* and *cht1* was analyzed at TP 12, 18, 24 and 36 hpi using RNAs extracted with column kits (A) or with the TRIzol reagent (B) from pools of 8 mosquitoes infected with *P. falciparum* from cage 1 and cage 2. The time points post infection (TP: 12, 18, 24 and 36 hpi) and the fold-change induction ($2^{-\Delta\Delta CT}$), using T12 as calibrator sample are shown. The $C_T$ values of the AgS7 housekeeping gene were used for normalization.

### Standard curves for absolute quantification by qPCR

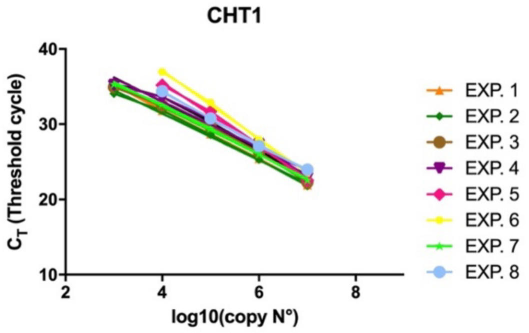

**CHT1 standard curves**

| EXP. | S | R2 | Y |
|---|---|---|---|
| 1 | -3,299 | 0,9996 | 45,13 |
| 2 | -3,098 | 0,9946 | 43,81 |
| 3 | -3,148 | 0,9904 | 44,88 |
| 4 | -3,075 | 0,9743 | 45,34 |
| 5 | -4,241 | 0,9958 | 52,48 |
| 6 | -4,524 | 0,9988 | 55,21 |
| 7 | -3,215 | 0,9973 | 45,27 |
| 8 | -3,495 | 0,9987 | 48,26 |

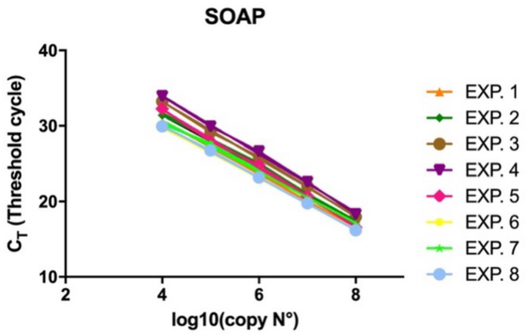

**SOAP standard curves**

| EXP. | S | R2 | Y |
|---|---|---|---|
| 1 | -3,551 | 0,9985 | 44,92 |
| 2 | -3,572 | 0,9969 | 46,01 |
| 3 | -3,774 | 0,9988 | 48,29 |
| 4 | -3,857 | 0,9984 | 49,41 |
| 5 | -3,860 | 0,9976 | 47,60 |
| 6 | -3,386 | 0,9996 | 43,41 |
| 7 | -3,382 | 0,9961 | 44,22 |
| 8 | -3,453 | 0,9996 | 43,88 |

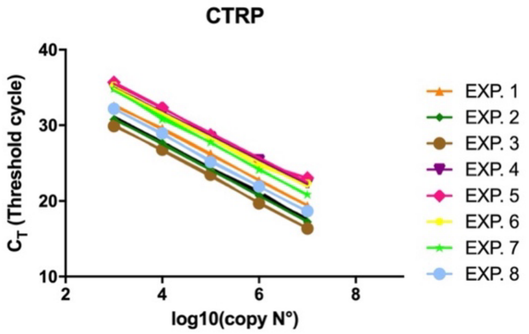

**CTRP standard curves**

| EXP. | S | R2 | Y |
|---|---|---|---|
| 1 | -3,326 | 0,9991 | 42,73 |
| 2 | -3,393 | 0,9996 | 41,06 |
| 3 | -3,414 | 0,9990 | 40,31 |
| 4 | -3,175 | 0,9990 | 44,47 |
| 5 | -3,258 | 0,9920 | 45,26 |
| 6 | -3,253 | 0,9929 | 44,64 |
| 7 | -3,463 | 0,9981 | 45,00 |
| 8 | -3,410 | 0,9993 | 42,43 |

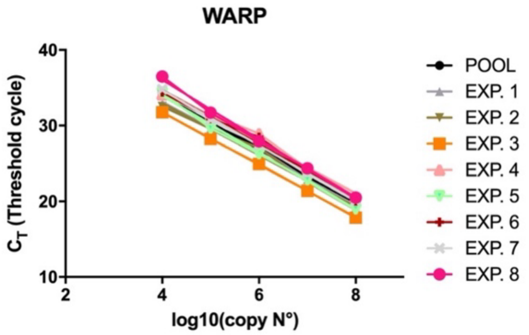

**WARP standard curves**

| EXP. | S | R2 | Y |
|---|---|---|---|
| 1 | -3,413 | 0,9940 | 46,79 |
| 2 | -3,438 | 0,9899 | 46,75 |
| 3 | -3,468 | 0,9998 | 45,64 |
| 4 | -3,418 | 0,9856 | 48,39 |
| 5 | -3,762 | 0,9963 | 48,99 |
| 6 | -3,569 | 0,9915 | 48,97 |
| 7 | -3,650 | 0,9994 | 49,28 |
| 8 | -3,931 | 0,9970 | 51,79 |

**Figure S2. Standard curves for the absolute quantification of ookinete markers through qPCR.** Standard curves obtained for each ookinete marker in 8 different experiments. Transcript copy number and corresponding threshold cycle ($C_T$) relative to the 5 serial dilutions points are reported on the X- and Y-axis, respectively. Slopes (S), correlation coefficients ($R^2$) and intercepts (Y) are reported in the tables on the right side.

#### Comparison among standard curves generated in independent experiments

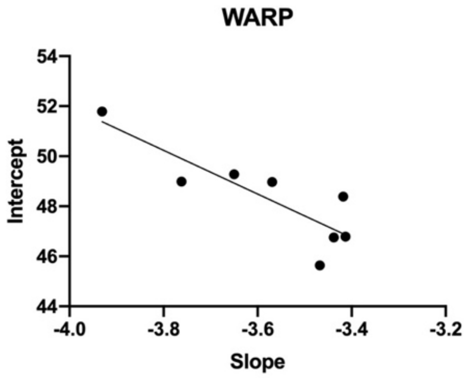

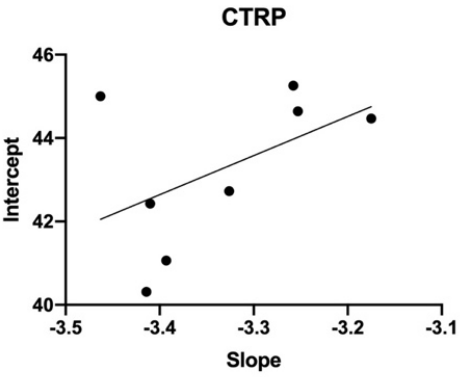

| | Slope vs. Intercept |
|---|---|
| Pearson r | |
| r | -0.8560 |
| 95% confidence interval | -0.9735 to -0.3814 |
| R squared | 0.7328 |
| P value | |
| P (two-tailed) | 0.0067 |
| P value summary | ** |
| Significant? (alpha = 0.05) | Yes |
| Number of XY Pairs | 8 |

| | Slope vs. Intercept |
|---|---|
| Pearson r | |
| r | 0.4970 |
| 95% confidence interval | -0.3196 to 0.8900 |
| R squared | 0.2470 |
| P value | |
| P (two-tailed) | 0.2102 |
| P value summary | ns |
| Significant? (alpha = 0.05) | No |
| Number of XY Pairs | 8 |

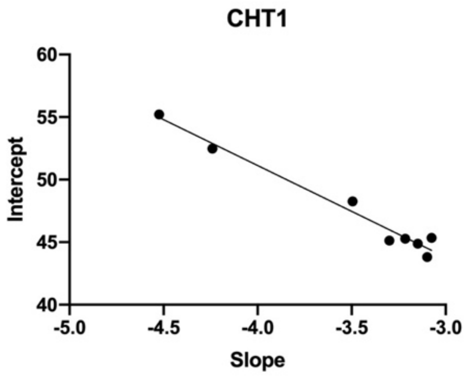

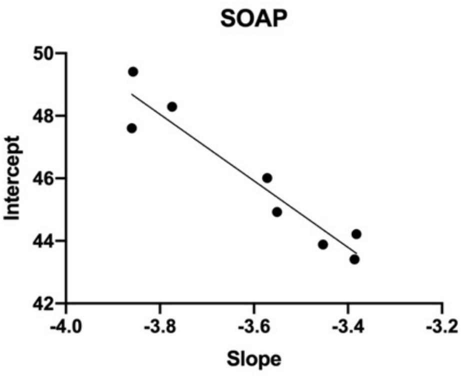

| | Slope vs. Intercept |
|---|---|
| Pearson r | |
| r | -0.9868 |
| 95% confidence interval | -0.9977 to -0.9262 |
| R squared | 0.9738 |
| P value | |
| P (two-tailed) | <0.0001 |
| P value summary | **** |
| Significant? (alpha = 0.05) | Yes |
| Number of XY Pairs | 8 |

| | Slope vs. Intercept |
|---|---|
| Pearson r | |
| r | -0.9552 |
| 95% confidence interval | -0.9921 to -0.7662 |
| R squared | 0.9124 |
| P value | |
| P (two-tailed) | 0.0002 |
| P value summary | *** |
| Significant? (alpha = 0.05) | Yes |
| Number of XY Pairs | 8 |

**Figure S3. Analysis of standard curves parameters.** Correlation between slope and intercept values of the four ookinete marker's standard curves in the 8 rounds of experiments. Results of Pearson's correlation test are reported in the tables below each graph.

#### Expression profile of the four ookinete markers in single mosquitoes

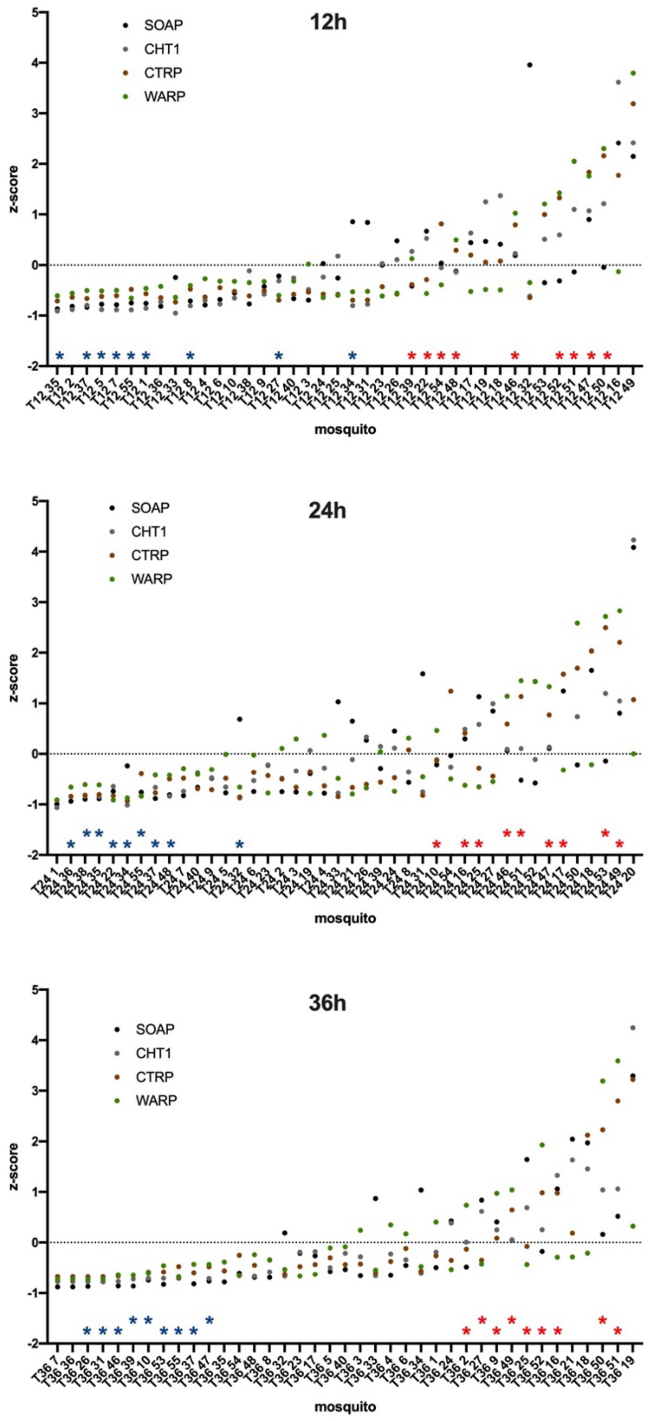

**Figure S4. Distribution of four ookinete marker expression across individual mosquitoes at different time points**. Scatter plots showing z-score–normalized transcript levels of soap, cht1, ctrp, and warp in individual mosquitoes at 12, 24, and 36 hpi. Each dot represents the standardized expression value of a given marker in a single mosquito. Mosquitoes are ordered along the x-axis according to increasing infection intensity (left to right). The horizontal dashed line indicates a z-score of 0 (mean expression level). Blue and red asterisks identify mosquitoes belonging to distinct infection intensity groups (e.g., low vs. high ookinete burden), as defined in the study. At every time point, increasing infection intensity is associated with progressively higher standardized expression of invasion-related markers, particularly cht1, ctrp, and warp, while soap shows greater inter-individual variability.

### Cluster analyses of RNA-seq samples

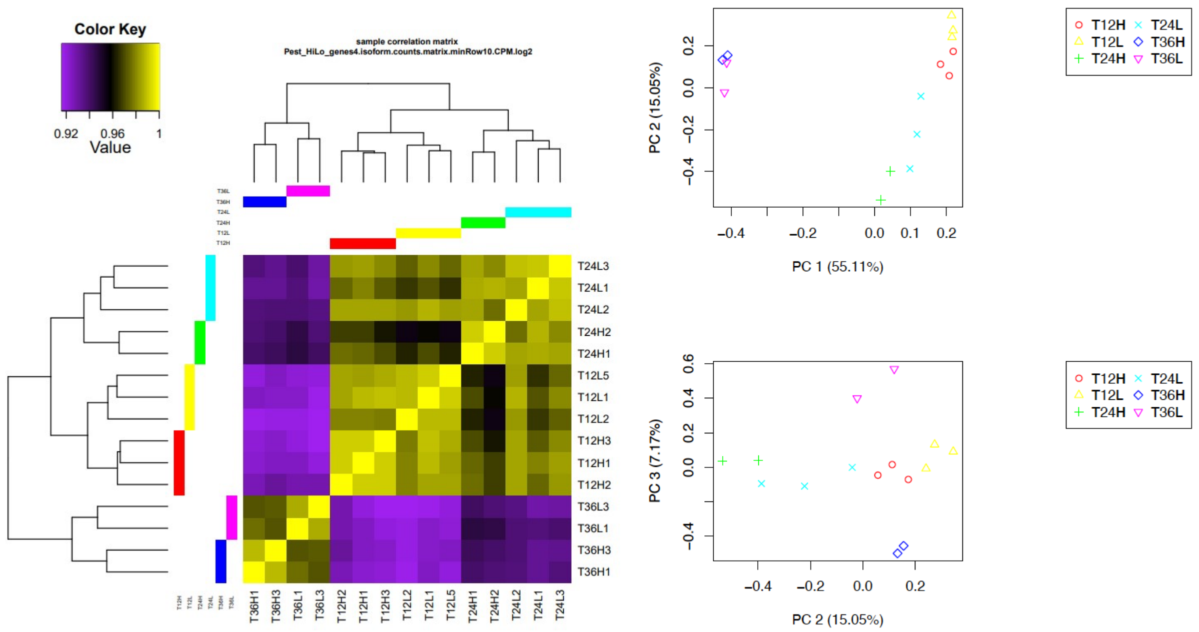

**Figure S5. Correlation matrix of the *An. coluzzii* infected with *P. falciparum* whole dataset.** Sample correlation matrix shows the overall similarity in expression profile for each sample at the three different time points post infection. At least two biological replicate per time points and for each condition were selected and analyzed. Yellow colour intensity indicates increasing sample correlation, whereas purple colour intensity indicates decreasing sample correlation. Dendrogram clustering on the X and Y-axis indicates the overall similarity between all samples. On the right panels, PCA analyses is reported.

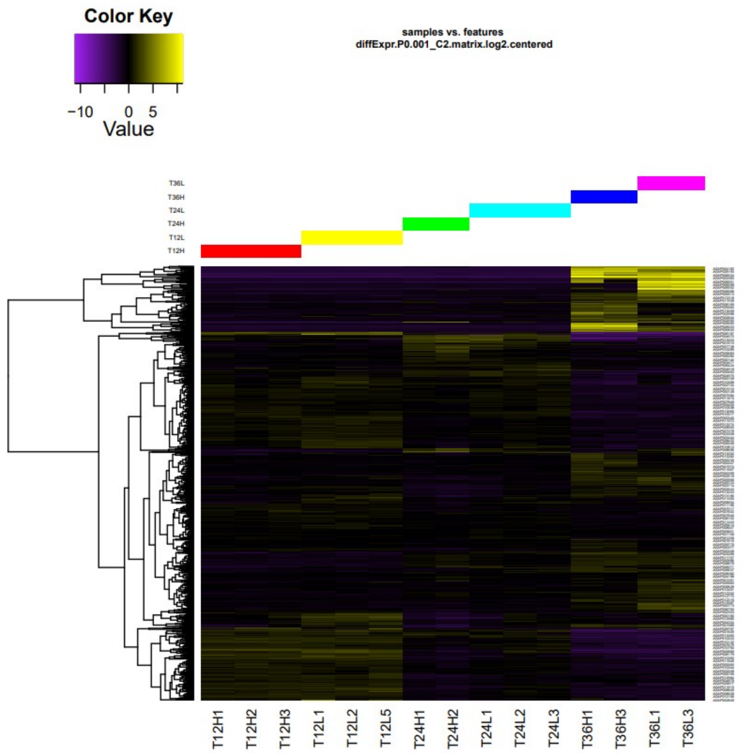

**Figure S6. Cluster analysis.** Differentially expressed (DE) genes of all sequenced samples were analysed and clusters created.

#### *Plasmodium* DE in RNA-seq databases

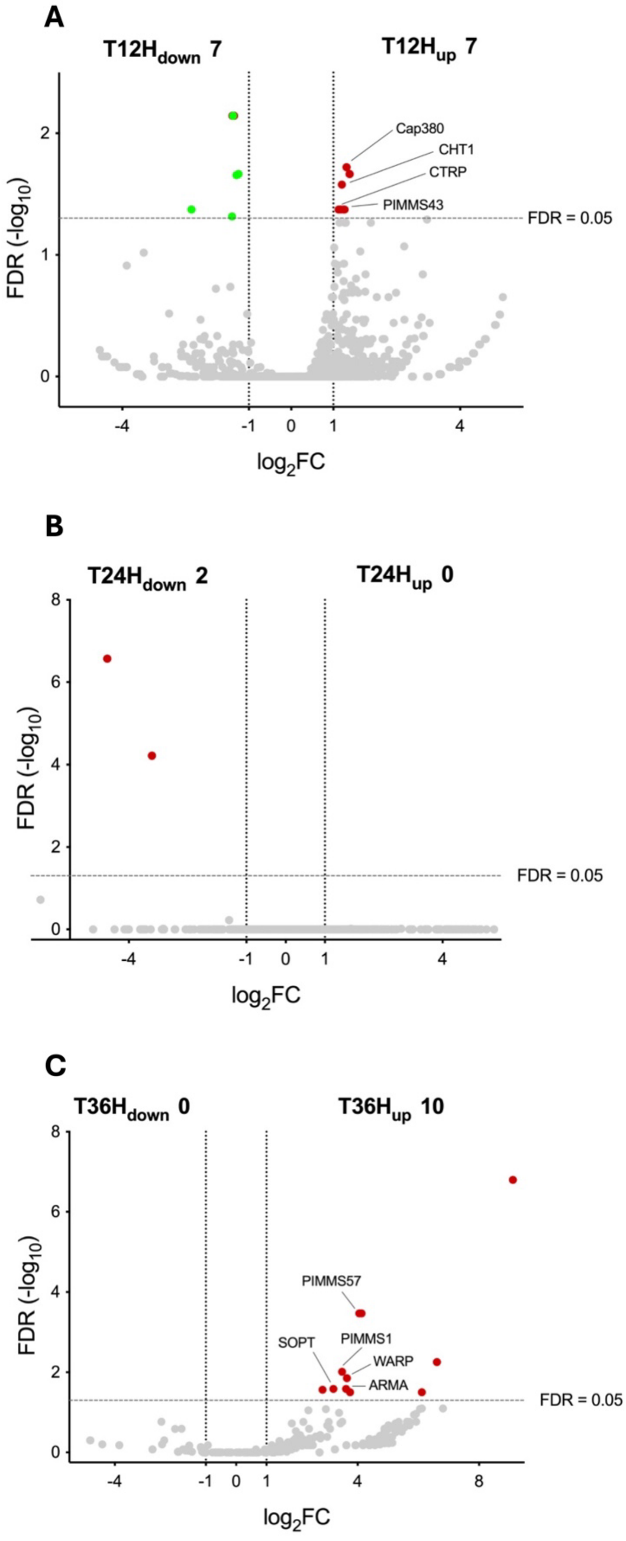

**Figure S7. Volcano plot showing *P. falciparum* DE genes at 12 (A), 24 (B) and 36 (C) hpi, comparing H vs. L datasets.** Reads from RNA-seq datasets were mapped using *P. falciparum* genome (3D7) dataset as reference. *Plasmodium* DE genes (FDR<0.05, -2>FC>+2) are highlighted in red (upregulated in H samples) or in green (upregulated in L samples).

#### Correlation among qPCR and RNA-seq values (markers' copies vs TPM)

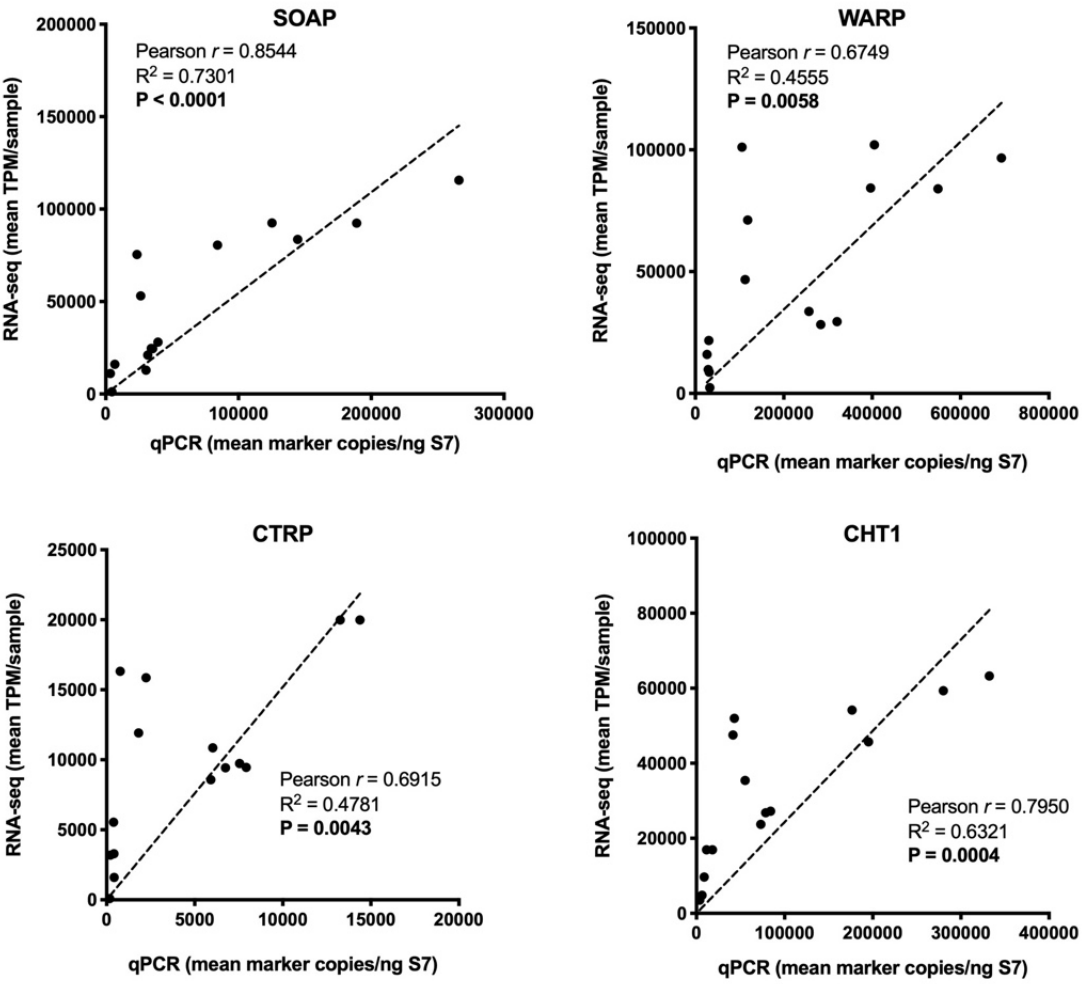

**Figure S8. Correlation of *P. falciparum* ookinete markers expression by qPCR and RNA-seq data.** Graphs show the correlation analysis of RNA-seq and qPCR data. Quantitative data derived from qPCR (expressed as marker copy number per ng of S7) and RNA-seq (expressed as marker TPM) for the four ookinete-specific markers are plotted along the x- and y-axes, respectively. The insets display the outcomes of Pearson correlation analyses. Each data point represents an individual sample.

### Gene ontology enrichment analysis at 36 hpi.

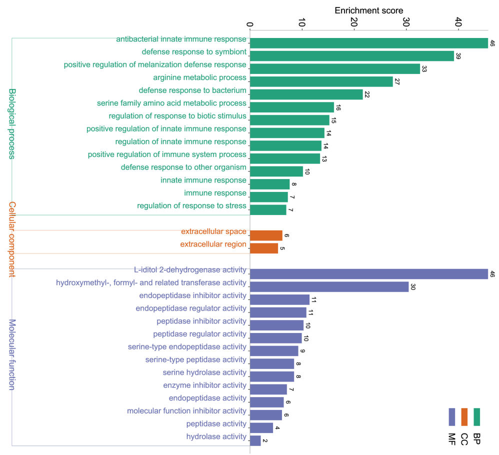

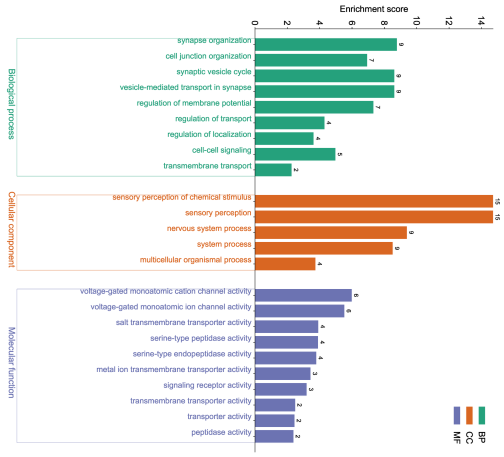

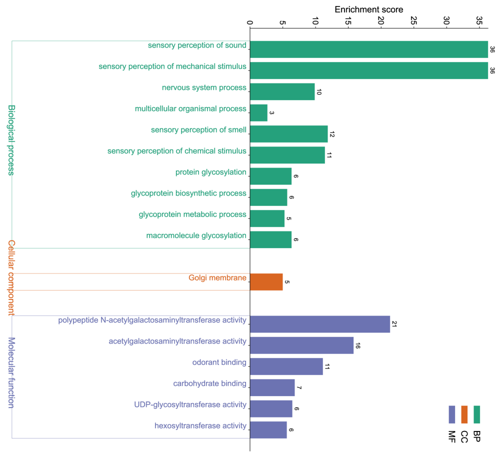

**Figure S9. GO enrichment analysis in mosquitoes with high versus low infection intensity at 12, 24 and 36 hpi.** Gene Ontology (GO) enrichment analysis of downregulated DE genes at T12 (A), T24 and upregulated at T36. Enriched GO terms are grouped by Biological Process (BP, green), Cellular Component (CC, orange), and Molecular Function (MF, purple). Bars represent enrichment scores, with corresponding numbers indicated (graphs were elaborated by Rplots).

**Table S1.** Summary of sequencing data.

| sample | Reads | total base pairs(bp) | filtered reads | mapped to PEST genome |
|---|---|---|---|---|
| T12 L1 | 43,524,334 | 12,375,825,439 | 43,486,236 | 33,208,454 |
| T12 L2 | 42,863,496 | 12,160,989,009 | 42,826,630 | 32,534,563 |
| T12 L5 | 39,993,022 | 11,283,220,365 | 39,950,291 | 29,707,165 |
| T12 H1 | 46,384,334 | 13,191,708,766 | 46,342,680 | 36,144,775 |
| T12 H2 | 47,000,033 | 13,377,791,717 | 46,964,423 | 36,151,519 |
| T12 H3 | 38,646,269 | 10,958,221,394 | 38,602,750 | 29,923,040 |
| T24 L1 | 37,239,060 | 10,492,757,611 | 37,202,155 | 27,953,571 |
| T24 L2 | 44,571,610 | 12,698,164,060 | 44,538,502 | 32,904,183 |
| T24 L3 | 37,178,373 | 10,523,659,845 | 37,145,136 | 27,522,038 |
| T24 H1 | 45,930,809 | 13,083,455,518 | 45,896,040 | 34,117,493 |
| T24 H2 | 52,500,356 | 14,861,288,953 | 52,455,969 | 38,430,429 |
| T36 L1 | 48,113,140 | 13,638,981,795 | 48,075,298 | 35,615,600 |
| T36 L3 | 43,543,117 | 12,306,927,331 | 43,502,448 | 32,333,474 |
| T36 H1 | 55,368,129 | 15,734,226,585 | 55,314,873 | 40,854,274 |
| T36 H3 | 41,145,797 | 11,601,523,756 | 41,110,166 | 30,151,656 |
| Total | 664,001,879 | 188,288,742,144 | 663,413,597 | 497,552,234 |
